# Petroleum Hydrocarbon Concentration as the Primary Driver of Soil Bacterial Community Dynamics During Mycorrhizal Fungi and Rhizobacteria Assisted Phytoremediation

**DOI:** 10.1101/2025.08.19.671157

**Authors:** Prama Roy, Barbara Zeeb

## Abstract

Petroleum hydrocarbon (PHC) contamination poses widespread environmental risks in Canada, where it is known to contribute to 60% of existing contaminated sites. It requires effective remediation strategies in the boreal ecozone due to the cold climate resulting in slow site weathering processes and soil nutrient cycling. This study utilized PHC-contaminated field soil (25,700 mg/kg total petroleum hydrocarbons - TPH) and background soil (<120 mg/kg TPH) collected from a Canadian boreal site where PHCs had weathered in place for 25+ years. Additionally, a 1:1 diluted soil was prepared (12,600 mg/kg TPH), and with the field contaminated soil, used to investigate the effects on rhizobacterial community composition. In two-year greenhouse experiments (2023 - 2025), the soils were planted with native or naturalized plant species and inoculated with i) arbuscular mycorrhizal fungi (AMF), ii) the biosurfactant-producing and plant-growth-promoting rhizobacterium (PGPR) - *Bacillus subtilis ATCC* 21332, or iii) both AMF and *B. subtilis* 21332. PHC contamination was determined to exert a dominant influence on rhizobacterial community composition. Alpha- and beta-diversity analyses revealed that neither plant species nor microbial inoculants significantly altered bacterial diversity or community structure beyond the overriding effect of PHC concentration. Proteobacteria, Actinobacteriota, Acidobacteriota, and Chloroflexi dominated across all soils, with hydrocarbon-degrading genera such as *KCM-B 112* and *Sphingomonas* significantly enriched in the 25,700 mg/kg TPH soil. Functional gene profiling identified widespread aerobic (e.g., *alkB*, *CYP153*, *assA*, *nahAc*, *pheA*, *xylM*, *xylE, todC1*, *bphA1,* and *pahE*) hydrocarbon-degradation pathways across phyla, suggesting extensive horizontal gene transfer and functional redundancy across treatments and plant species. Exogenously introduced *B. subtilis* ATCC 21132 established only in the background soil, suggesting competitive dominance of indigenous PHC-degraders. These findings underscore the magnitude of PHC concentration as the primary driver of rhizobacterial dynamics and indicate that augmenting native microbial capacity through soil carbon enhancement, rather than bioaugmentation, may significantly impact soil bacterial community composition during rhizodegradation of boreal soils.

## 1.0 Introduction

Petroleum hydrocarbon (PHC) contamination of soil, resulting from extensive extraction, refining, transport, and storage of petroleum products, represents one of the most pressing environmental challenges globally. The widespread occurrence of petroleum spills and leakage from storage facilities has led to substantial contamination of terrestrial and aquatic ecosystems, posing significant risks to human health and environmental integrity (Tang et al, 2011; Khan et al., 2018). The boreal ecozone is the largest in Canada, encompassing >50% of Canada’s total land mass (CCME, 2008c). Approximately 60% of Canadian contaminated sites are polluted with PHCs. Traditional remediation approaches, including excavation and *ex-situ* treatment, are often prohibitively expensive and environmentally disruptive, necessitating the development of sustainable, cost-effective *in-situ* bioremediation strategies (Hale et al., 2021). Bioremediation research in the boreal ecozone is needed as a limited number of plants and microbes are adapted to the cold climate, slow nutrient cycling, and podzolic soils characterized by limited nitrogen and phosphorous (Giesler et al., 2002; Deluca and Boisvenue, 2012; Sponseller et al., 2016).

Rhizodegradation, also known as rhizoremediation, has emerged as a particularly promising approach for the sustainable ‘green’ remediation of PHC-contaminated soils (Correa-García et al., 2018). This bioremediation biotechnology leverages the synergistic relationship between plants and their associated rhizosphere microorganisms to enhance the degradation of organic contaminants (Hussain et al., 2018). The fundamental principle underlying rhizodegradation is the ‘rhizosphere effect’, whereby plant root exudates actively modify their surrounding soil environment through the exudation of diverse organic compounds, creating a unique ecological niche that supports enhanced microbial activity and contaminant degradation (Cheng et al., 2019). Plant roots release a complex mixture of exudates, including organic acids, amino acids, sugars, terpenoids, and secondary metabolites, which serve multiple functions in contaminant remediation (Eze and Amuji, 2024). These compounds provide carbon and nutrient sources for PHC-degrading bacteria, enhance the bioavailability of PHCs through desorption mechanisms, and create favorable microenvironmental conditions for bacterial proliferation (Liao et al., 2021). Furthermore, root exudates can directly participate in hydrocarbon transformation through enzymatic processes involving the action of plant-derived peroxidases, laccases, and phenol oxidases (Rohrbacher and St-Arnaud, 2016).

Arbuscular mycorrhizal fungi (AMF) represent a critical component of the rhizosphere microbiome that can greatly enhance rhizodegradation efficiency (Rabie, 2005; Ren et al., 2017). AMF form symbiotic associations with approximately 80-90% of terrestrial plant species, creating extensive hyphal networks that expand the effective rhizosphere and increase plant access to nutrients and contaminants. The presence of AMF has been shown to improve PHC degradation through multiple mechanisms, including enhanced plant stress tolerance in contaminated environments, increased soil enzyme activity (peroxidases, laccases, catalases), and improved bioavailability of PHCs (Ingrid et al., 2016; Rajtor and Piotrowska-Seget, 2016).

Bioaugmentation strategies involving the introduction of plant growth-promoting rhizobacteria (PGPR) with hydrocarbon-degrading capabilities represent another promising approach for enhancing rhizodegradation (Hou et al., 2015; Sampaio et al., 2019). *Bacillus subtilis* is recognized for its dual capacity for plant growth promotion and aliphatic PHC biodegradation through secretion of alkane hydroxylases and alcohol dehydrogenases (Parthipan et al., 2017). This rhizobacterium naturally produces lipopeptide biosurfactants (primarily surfactin), which reduce surface tension to enhance hydraulic conductivity of soils and the bioavailability of PHCs (Ahimou et al., 2000). The surfactin produced by *B. subtilis* ATCC 21332 is especially potent, reducing surface tension of water from 72 mN/m to as low as 27 mN/m at room temperature (Cooper et al., 1981; Fox and Bala, 2000), outperforming synthetic surfactants like sodium dodecyl sulfate and cetyltrimethylammonium bromide (Sakthipriya et al., 2021).

This study addresses critical knowledge gaps in the literature regarding the combined effectiveness of AMF and PGPR at high (>10,000 mg/kg) PHC contamination levels by investigating the rhizodegradation of PHCs using plants in combination with AMF and *Bacillus subtilis* bioaugmentation in 25,700 mg/kg total petroleum hydrocarbon (TPH) and/or 12,600 mg/kg TPH soil (Li et al., 2023; Hnini et al., 2024). Petroleum-tolerant, perennial, regionally native or naturalized plants were prioritized to ensure both effective rhizodegradation and ecological compatibility (Aprill and Sims, 1990; Olson et al., 2007; Reynoso-Cuevas et al., 2011; Papik et al., 2023). *Andropogon gerardii* (big bluestem), *Bouteloua curtipendula* (sideoats grama), *Panicum virgatum* (switchgrass) are three tufted C4 prairie grasses whose deep, fibrous root systems oxygenate the rhizosphere and have accelerated remediation of soils containing 10,000 to 25,000 mg/kg TPH (Thomas et al., 2013; McIntosh et al. 2016). *Picea mariana* (black spruce), the keystone conifer of the eastern boreal forest, maintains ≥75% germination and robust seedling growth in weathered soils with 11,900 mg/kg TPH, making it a suitable candidate for long-term site reclamation (Roy et al., 2023). *Achillea millefolium* (common yarrow) adds functional diversity as a hardy forb whose aromatic, finely branched roots remain vigorous in soils containing 10,000 to 50,000 mg/kg TPH and have recorded 45–65% TPH reduction in pot trials (Masu et al., 2014).

This research is the first to examine the dual symbioses of AMF and PGPR bioaugmentation in boreal site soils. This study focuses particularly on characterizing soil bacterial community responses to different treatment combinations and PHC contamination levels, providing insights into the mechanisms underlying enhanced PHC rhizodegradation in plant-microbe-fungi systems. The findings contribute to the development of sustainable, cost-effective bioremediation technologies that can be used to optimize rhizodegradation in PHC-contaminated boreal environments. Understanding the complex interactions between indigenous soil bacteria, introduced PGPR, and plant hosts is essential for developing robust, field-applicable rhizodegradation strategies (Dong et al., 2014).

## 2.0 Methods

### 2.1 Site Soils

#### 2.11 Soil Collection

In August 2022, background (120 mg/kg total petroleum hydrocarbons) and contaminated (25,700 mg/kg TPH) site soil was collected from the site of a former petroleum distribution terminal located in northern Ontario, where frequent oil spills occurred (Roy et al., 2025). The site held No. 2 diesel fuel and has been inactive for 25+ years, indicating significant weathering of the soils. No refinery processes occurred at the site. The contaminated site was previously sampled for soils in earlier toxicological studies carried out by Roy et al. (2023, 2024), which revealed high toxicity to native plants and soil invertebrates in the 11,900 – 12,800 mg/kg TPH concentration range. However, the contaminated soil in this study was collected from a different hotspot region, located approximately 80 m away from the previously studied contaminated region. The background soil in this study was collected from the same location used for background soil collection in our earlier studies (Roy et al. 2023, 2024). Both background and contaminated soils were collected from 6 m^2^ plots using an excavator, to a depth of 60 cm. The top 10 cm of soil was excluded to remove plant biomass. A total of 42 barrels (200-L) were filled with soil (21 barrels for each soil) and subsequently transported to the Royal Military College of Canada (RMC) for this greenhouse study.

#### 2.12 Soil Preparation

The <120 mg/kg TPH and 25,700 mg/kg TPH soils were sieved through a 6.25 mm soil sifter box, and large rocks were manually removed. Subsequently, the soils were potted into DCN Harmony Planters (30.5 cm length x 30.5 cm width x 45.6 cm height) to a volume of 25 L per pot. Due to the high PHC concentration in the contaminated site soil (Table 2), a 1:1 diluted mixture of the <120 mg/kg TPH: 25,700 mg/kg TPH site soils was created to test phytoremediation at a lower PHC concentration (12,600 mg/kg TPH ‘diluted soil’). Each pot was filled with an equal volume of background and contaminated soil and mixed manually until a homogenous texture/color was achieved. All pots were sealed with aluminum foil and heavy-duty duct tape, and stored in a cool location to limit volatilization of PHCs prior to the start of the experiments.

#### 2.13 Soil Particle Size and Nutrients

Particle size of the soils were measured at ALS Environmental using the ASTM D6913 and D7928 reference methods (ASTM 2014, 2021). As >50% of the soils have a grain diameter of > 0.075 mm, the <120 mg/kg TPH, 25,700 mg/kg TPH, and 12,600 mg/kg TPH soils are all considered coarse-grained (Supplementary Information Table S1; CCME, 2008a). At the start of the greenhouse experiments, soil nutrient analyses were performed by the University of Guelph Laboratory Services (Table S2), as described by Roy et al. (2023, 2024). One method blank, certified reference material, and duplicate was analyzed with the soils, to the standard of ISO/IEC 17025 (ISO, 2005). The relative percent difference between duplicate samples was <20% for all parameters. The <120 mg/kg TPH soil contains 4x, 7x, 2x, 1.5x, and 1.5x higher nitrogen from ammonium (Ammonium-N), nitrate-N, nitrite-N, potassium, and phosphorous, respectively, compared to the 25,700 mg/kg TPH soil (Table S2). The 25,700 mg/kg TPH soil contains 5x higher total carbon and organic carbon, and 6x higher inorganic carbon than the <120 mg/kg TPH soil. The <120 mg/kg TPH soil is basic (pH 7.4) whereas the 25,700 mg/kg TPH is more acidic (pH 6.4).

#### 2.14 Initial Soil Contaminant Concentrations

In December 2022, the Canadian Council of Ministers of the Environment (CCME, 2008b) Fractions 2-4 concentrations in the <120 mg/kg TPH and 25,700 mg/kg TPH soils were measured at the CALA-accredited Analytical Services Unit at Queen’s University from three randomly-selected barrels, for each soil (F2: >C_10_-C_16_, F3: >C_16_-C_32_; F4: >C_32_ -C_50_ effective carbon numbers; Table 2). Three soil samples were analyzed from each barrel using 1:1 hexane: acetone solvent extraction, ultrasonication, rotary evaporation (Büchi rotovapor R-114), silica gel cleanup, and gas-chromatography with flame ionization detection (Agilent GC 7890A), based on the CCME (2008c) reference method. In July 2023, the CCME F2-F4 concentrations in the diluted soil were sampled from six unplanted greenhouse pots using the same analytical methods. One method blank, one sample duplicate, and a 5,000 mg/kg diesel fuel quality control spike was used in each soil PHC analysis.

The <120 mg/kg TPH and 25,700 mg/kg TPH soils were also screened for metal contaminants at the Analytical Services Unit using the US EPA 200.7 method for reference (US EPA, 1994). Soil samples (0.5 g each) were ground up and dried overnight and then subject to a modified *aqua regia* digestion (7 mL water, 6 mL HCl, and 2 mL HNO_3_) on a DigiPREP LS graphite block (SCP Science) for four hours at 95 °C. The digested samples were subsequently analyzed using inductively-coupled plasma-optical emission spectroscopy (Thermo iCAP 7400 Dual view spectrometer). One certified reference material, one duplicate, and one method blank was also analyzed. Neither soil was contaminated with metals (Table S3).

### 2.2 Rhizodegradation Experiments

#### 2.21 Plant Species

Seeds of *P. mariana* were shipped to the RMC from the Saskatchewan Research Council in 2015. Seeds of *A. gerardii, B. curtipendula*, *A. millefolium*, and *P. virgatum* were shipped from Sheffield’s Seed Company in 2022 (lot numbers: 1834306, 1832814, 1833869 and 1834252). Germination tests for each species were conducted in parafilm-sealed Petri dishes filled with 10 g of background site soil and ten seeds that were moistened with deionized water. The Petri dishes were left in the dark at room temperature for two weeks; all Petri dishes showed >80% germination.

#### 2.22 Experimental Design

In January 2023, a two-year greenhouse phytoremediation study was initiated at RMC using the <120 mg/kg TPH and 25,700 mg/kg TPH soils (Figure 1). The five selected plant species and four experimental treatments were employed to evaluate the most successful treatment for phytoremediation. The first treatment was a ‘Plants-Only Control’ with no soil amendments. The second treatment (‘AMF’) involved inoculating each pot with 13.2 g of with MycoApply Ultra Fine Endo ©. This commercial blend is comprised of 130,000 propagules/lb. of four Glomeromycota arbuscular mycorrhizal fungi: *Rhizophagus irregularis* (formerly *Glomus intraradices*)*, Glomus mosseae, G. aggregatum,* and *G. etunicatum*. Four of the plant test species, excluding *P. mariana*, have previously shown increased growth after forming symbiotic mycorrhizae with AMF of the *Glomus/Rhizophagus* genus (Roudi and Salamatmanesh, 1998; Gustafson and Casper, 2006; Reinhart and Anacker, 2014; Bindell et al., 2021).

**Figure 1.**
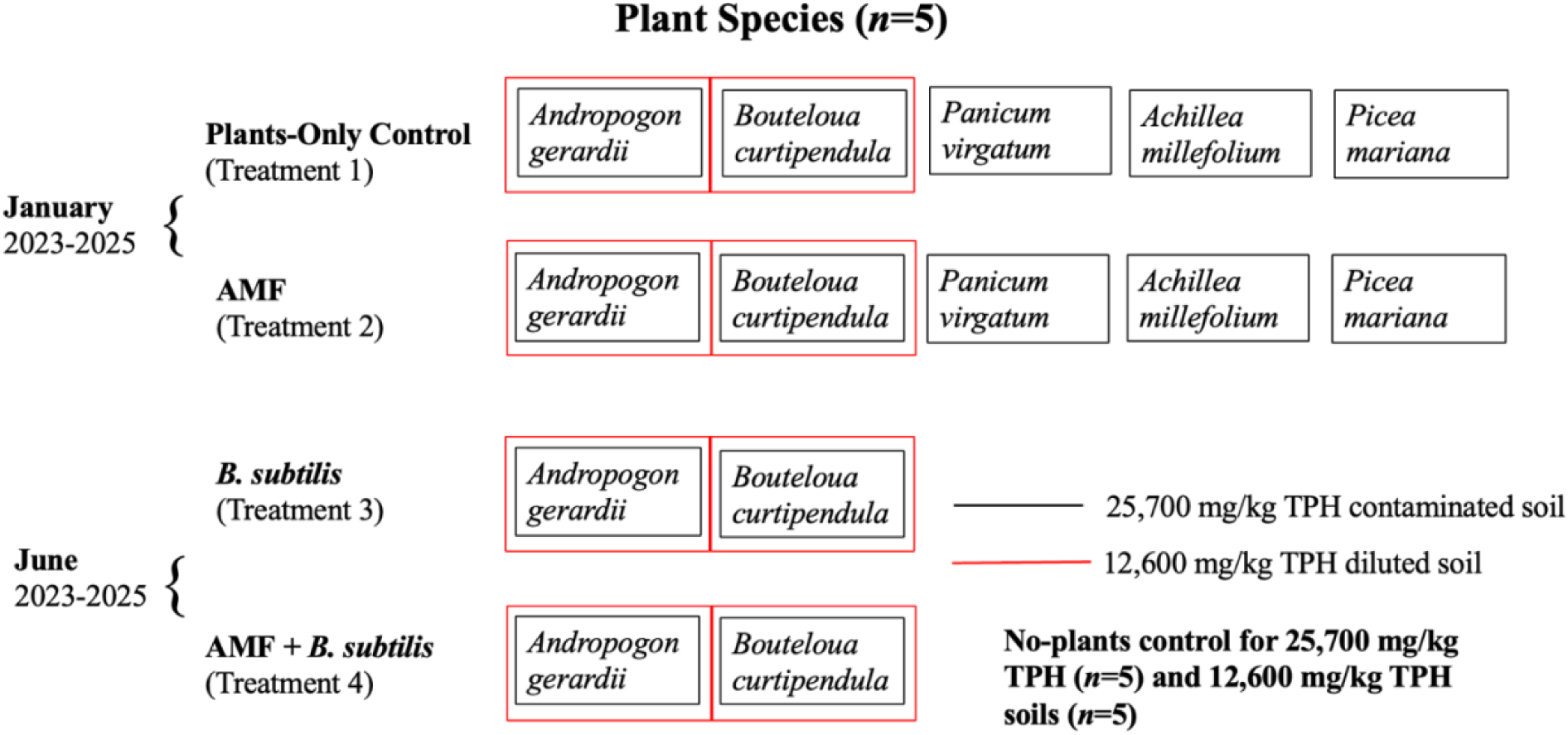
Experimental design of the two-year greenhouse phytoremediation experiments. Four treatments were employed: a Plants-Only Control (1), AMF (2), *B. subtilis* (3) and AMF + *B. subtilis* soil inoculants (4). Five plant species underwent treatments 1 and 2 in the <120 mg/kg TPH (background) and 25,700 mg/kg TPH (contaminated) soil from January 2023 to January 2025. *Andropogon gerardii* and *Bouteloua curtipendula* were subject to all treatments in the <120 mg/kg TPH and 12,600 mg/kg TPH (diluted) soil from June 2023 to June 2025. A no-plants control was also employed. Five replicates (*n*=5) were used for each treatment.

In June 2023, the two best-growing plant species (*Andropogon gerardii* and *Bouteloua curtipendula*) were planted in the <120 mg/kg TPH and 12,600 mg/kg TPH soils (Figure 1). The Plants-Only Control and AMF treatments were utilized. A third treatment (‘*B. subtilis*’) involved inoculating the soils with *Bacillus subtilis* ATCC 21332, an endospore-forming, non-pathogenic and plant-growth-promoting-rhizobacterium (PGPR). This strain of *B. subtilis* also produces biosurfactants (surfactin, iturin, fengycin). A total of 42 mL of active *B. subtilis* ATCC 21332 culture was added to each treatment pot (average absorbance at 600 nm (OD_600_): 1.87; cell dry weight: 4 g/L). A fourth treatment (‘AMF + *B. subtilis*’) involved inoculating soils with the same volume of AMF and *B. subtilis*.

#### 2.23 Seeding Rate

Before planting, *P. mariana* seeds were cold-stratified in *Sphagnum* peat moss and deionized water for six weeks to improve germination. Seeds of the other four species were hydrated for 24 hours for scarification. Seeds were evenly sown into the top 1 cm of soil. For all species except *P. mariana*, the seeding rates that were used for the treatments that started in January 2023 were 20% higher than the pure live seed counts for each seed cultivar (Table 1). Twelve *P. mariana* seeds per pot were sown, based on the seed lot’s high germination rate during earlier toxicity testing (Roy et al., 2023).

**Table 1.** Seeding rates for four of the plant species used during the greenhouse experiments. Pure Live Seed calculations for each cultivar were provided by the United States Department of Agriculture (USDA).

| Plant Species | Pure Live Seed (lb/acre) | Seed Density Preference (seeds/ft<sup>2</sup>) | Seeds Added (January 2023) | Seeds Added (June 2023) | Reference |
| --- | --- | --- | --- | --- | --- |
| *Andropogon gerardii* | 9 | 34 | 41 | 41 | USDA, 2015 |
| *Bouteloua curtipendula* | 3 | 13 | 16 | 16 | USDA, 2014 |
| *Panicum virgatum* | 5 | 44 | 53 | -- | USDA, 1997 |
| *Achillea millefolium* | 2 | 51 | 61 | -- | USDA, 2022 |

#### 2.24 Greenhouse Conditions

The greenhouse at RMC was set to a temperature range of 20°C (winter) to 30°C (summer) using an iGrow 1800 Advanced Environmental Controller. The LinkConn 1000 v3.6 program was used. During the summer and winter months, the light intensity ranged from 260 to 660 W/m^2^ and 780 to 1120 W/m^2^, respectively. Natural sunlight was used unless light intensity dropped below 350 W/m^2^, where Verilux Full Spectrum H588 light rods were employed. Relative humidity was set to 50% throughout the year. Greenhouse pots were watered three days per week. Approximately 3 L of water was added to each pot on a weekly basis. Any weeds and non-target plant species were manually removed from the pots.

### 2.3 *Bacillus subtilis* ATCC 21332 Culturing

#### 2.31 Seed Cultures

In January 2024, a seed culture of *B. subtilis* ATCC 21332 (Cedarlane Labs) was created in a Corning baffled culture flask (500-mL) that was filled to 100 mL with Oxoid CM001 Nutrient Broth (Thermo Fisher). The culture was shaken at 230 rpm and 37 °C for 16 hours within a Thermo Fisher MaxQ incubated orbital shaker. Sterile sponges were used as flask covers to facilitate aeration of the culture. Purity of the culture was established by streaking the culture in a nutrient broth agar plate and verifying the sole presence of *B. subtilis* with oil immersion microscopy at 1000x magnification (Celestron Labs CB2000C microscope with WF10X ocular lens and 100X/1.25 oil objective lens).

Growth curves were created using UV-Vis spectrophotometry (Agilent Cary 60) and cell dry weight calculation (Figure S1A). *B. subtilis* was able to proliferate significantly in the seed culture over a 16-hour period. UV-Vis was carried out at an absorbance of 600 nm (OD_600_) within the linear range of the spectrophotometer (maximum linear absorbance = 3). A sterile borosilicate glass cell filtration unit, equipped with a stainless steel clamp and sterilized polyvinyl chloride 0.22-micron filter paper (Millipore Sigma), was used for calculating the cell dry weight. Every 2-3 hours, 5 – mL of culture was used for both cell dry weight and absorbance readings. Three absorbance readings were taken and averaged. For cell dry weight, an equal volume of sterile water was added to rinse away the nutrient broth. The pre-weighed filter papers were dried for 24 hours at 70 °C before weighing. The initial and final pH and surface tension of the seed culture was also measured using Hydrion pH paper and Precision Dunouy Tensiometer (CSC Scientific). A final OD_600_ of 1.91 and cell dry weigh of 0.89 g/mL was achieved.

#### 2.32 Mineral Salt Medium Cultures

Five (5-) mL of the 14-hour seed culture was used to inoculate 95-mL of sterile MSM in the 500-mL Corning baffled culture flasks. The use of MSM facilitates greater biosurfactant production than nutrient broth (Cooper et al., 1981; Tabatabaee et al., 2005). The MSM ingredients used are based on the formula used by Willenbacher et al. (2015). While glucose was used as the carbon source by Willenbacher et al. (2015), sucrose was used in this study because it is a common plant root exudate. Sucrose has been shown to facilitate greater *Bacillus* growth and surfactin production than glucose, despite requiring additional enzymes (i.e, sucrase and invertase) for metabolism, which likely evolved as a competitive species-level adaptation for carbon utilization in the rhizosphere (Liu et al., 2023).

The MSM culture was shaken at 230 rpm and 37 °C for 30 hours to create absorbance (Agilent Cary 60) and cell dry weight growth curves (Figure S1B). Ten (10-) mL of culture was used to determine cell dry weight. Absorbance and cell dry weight readings were taken every 2 - 10 hours. The initial and final pH and surface tension of the MSM culture was measured using Hydrion pH paper and Precision Dunouy Tensiometer (CSC Scientific). The 30-hour *B. subtilis* MSM cultures supplemented with 28.3 g/L sucrose was used to inoculate *B. subtilis* and the AMF + *B. subtilis* treatment pots (OD_600_ = 2.66 to 2.92).

#### 2.33 Sucrose Optimization

Colorometric reducing-sugar assays were conducted to optimize sucrose concentration in MSM culture (OD_600_ = 1.91) with 1% v/v of 3,5-dinitrosalicylic acid (DNS) reagent. The DNS was purchased from Millipore Sigma (CAS #609-99-4). The DNS reagent ingredients (w/v) used were taken from Miller (1959) and methods were optimized by Wang (n.d.) for HCl-mediated hydrolysis of sucrose into glucose and fructose. Four concentrations were tested in MSM culture: 9.4, 18.9, 28.3, and 37.7 g/L of sucrose. Absorbance was measured at 540 nm (OD_540_) on the Agilent Cary 60 spectrophotometer. Following the completion of the assays, insignificant (<5 g/L) quantities of sucrose remained in each tube, indicating complete utilization of sucrose by *B. subtilis* (Figure S2). *B. subtilis* were subsequently sub-cultured from the seed inoculum (5-mL) in 95 mL of sterile MSM to determine which sucrose concentration yielded the highest growth (230 rpm at 37 °C for 30 hours). *B. subtilis* growth increased with sucrose concentration until the 28.3 g/L concentration was reached (OD_600_= 2.69). As such, 28.3 g/L sucrose was utilized for culturing *B. subtilis* in MSM for the *B. subtilis* and AMF + *B. subtilis* treatments.

#### 2.34 Most Probable Number Assays

Most probable number (MPN) assays were used to enumerate viable cell counts of the MSM culture with sucrose (OD_600_ = 2.68). An MSM culture supplemented with 18.22 mL/L hexadecane (OD_600_ = 0.11; Millipore Sigma Cas #544-76-3) was grown under the same conditions as the MSM with sucrose and used to estimate whether *B. subtilis* can grow with a PHC as the sole carbon source. The hexadecane concentration (18.22 mL/L) was calculated based on the equivalent carbon number of 28.3 g/L sucrose in the MSM cultures.

A novel method for MPN assays, modified from Haines et al. (1996) and Van Hamme et al. (2000), involved performing a 10-fold serial dilution of the MSM cultures on Corning Clear Polystyrene 96-well microplates (Figure S3). The first row of each microplates was filled with 100 µL of the MSM culture supplemented with either sucrose or hexadecane. Sterile MSM with 28.3 g/L sucrose was added to rows 2-12. For the hexadecane plate, sterile MSM with 18.22 mL/L hexadecane was added to rows 2-12 individually due to emulsification. A serial dilution was performed by pipetting 10 µL of culture from rows 1 to 11, with row 12 remaining as the sterile control. Subsequently, 5 µL of filter-sterilized 150 mg/mL 2,6-dichlorophenolindophenol (DCPIP) was added to rows 2-12 as the redox dye (Millipore Sigma Cas #1266615-56-8). The microplates were aseptically sealed and then incubated at 37°C for 48 hours. Wells containing *B. subtilis* changed from blue to clear after the 48 hours. The number of colony forming units (CFU/mL) was estimated using the ‘MPN’ package in R. The MPN assay of the *B. subtilis* MSM culture with 28.3 g/L sucrose showed bacterial growth corresponding to 8.31 x 10^8^ CFU/mL of *B. subtilis* (Figure S3). The MPN assay of the *B. subtilis* MSM culture with 18.22 mL/L hexadecane grew to 252 CFU/mL *B. subtilis*. Row 12 of both assay plates remained blue, confirming no contamination of the negative control.

#### 2.35 Hemocytometry

Hemocytometry (Hausser Scientific Improved Neubauer) counts were performed to enumerate living cells by diluting samples of the fully grown seed (OD_600_ = 1.78) and MSM cultures (OD_600_ = 2.71) with trypan blue dye, using the conventional method outlined in Zhang et al. (2019). The seed culture was stained at a concentration of 6:1 trypan blue to culture while the MSM cultures were stained 48:1 with trypan blue. The hexadecane culture was stained at a ratio of 1.2x trypan blue to culture. Each count was performed in triplicate, and the averages were used to compare cellular growth across the different media. The average number of *B. subtilis* cells in the seed inoculum and MSM with sucrose were 3.0 x 10^8^ cells/mL and 2.50 x 10^9^ cells/mL, respectively. A total of 1.26 x10^6^ cells/mL was found in the MSM with hexadecane culture (OD_600_ = 0.11).

#### 2.35 Critical Micelle Concentration

The critical micelle concentration (CMC) of the MSM culture with sucrose (OD_600_ = 2.73) was calculated using the Precision Dunouy Tensiometer to determine the efficiency of biosurfactant production by the culture, using the method described in Sheppard and Mulligan (1987). The CMC of the *B. subtilis* MSM culture supplemented with 28.3 g/L sucrose was achieved at a concentration of 1.6% v/v culture in sterile water where a surface tension of 31 mN/m was achieved at 23°C (Figure S4). The low culture concentration at which the CMC wasachieved is indicative of highly efficient extracellular biosurfactant production (Gudina et al., 2010; Perinelli et al., 2020). The surface tension of the culture did not decrease below 26 mN/m, which was achieved at 12.3% v/v culture in sterile water. The surface tension of the culture did not change with centrifugation at 8,000 rpm for 10 minutes, indicating that the biosurfactants produced by the *B. subtilis* strain are extracellular.

### 2.4 16S Metagenomics

#### 2.41 Soil Sampling

Soil samples (2 g) were isolated from the rhizosphere of each of the pots using sterile scoopulas following one year of plant growth from seed in the RMC greenhouse. The samples were collected from a depth of 5-60 mm, at a 10 mm distance from the plant roots, where bacteria are typically concentrated (Fierer et al., 2003; Ranjard et al., 2003; Song et al., 2022). The soil samples were stored in sterile 15-mL conical tubes and then refrigerated (4 °C) for one month prior to soil DNA isolation.

#### 2.42 DNA Isolation and Qubit Assays

In pots with plants, DNA was isolated from the rhizosphere soil samples using the Quick DNA Fecal-Soil Miniprep Kit from Zymo Research (catalog # D6010 from VWR International). Beta-mercaptoethanol (0.5% v/v in Genomic Lysis Buffer), purchased from Fisher Scientific (CAS #60-24-2), was used to denature proteins and improve cell lysis. An MP Fastprep-24 with 2-mL tube assembly was used for bead beating, according to the kit protocol. The filtered DNA was refrigerated (4 °C) for two weeks before library preparation. The Qubit dsDNA HS kit (Fisher Scientific Catalog #Q33231) and Qubit 2.0 fluorometer were used to quantify the total double-stranded DNA following extraction.

#### 2.43 DNA Library Preparation

The 16S SSU rRNA gene library was prepared using the Quick-16S Plus NGS Library Prep Kit from Zymo Research (Catalog # D6430-PS1). The V4 region of the 16S SSU rRNA gene was amplified using 515F/806R primers for a total amplicon size of 388 base pairs. Quantitative (q) PCR was carried out using the kit-recommended program on a CFX Opus 96 Real-Time PCR System.

The ZymoBIOMICs Microbial Community DNA Standard (2000 ng) was diluted with molecular-grade water into a series of standards (1 to 75 ng/µL) that were used to quantify bacterial DNA in each soil sample (catalog #D6306). The log of the quantity of DNA (ng/ µL) was plotted against critical thresholds reported from the thermal cycler and was used to create a linear standard curve with a Pearson correlation coefficient of 0.97 (Figure S5). The equation of the line from the standard curve (y = -8.7178x + 36.412) and reported critical threshold values from the qPCR run were used to enumerate the bacteria DNA before sequencing (Figures S6 - S7).

Unique dual index barcodes were tagged to each DNA library during the qPCR run. Two of the five amplified DNA libraries were selected based on highest DNA concentration. The DNA libraries were pooled together (6 µL each), and subsequently purified with magnetic beads. Samples were stored at 4 °C for one week and then sequenced.

#### 2.44 Next Generation Sequencing

The DNA in the pooled library was calculated using the Qubit dsDNA HS kit three times and subsequently averaged on the day of sequencing. The 16S Metagenomic Sequencing Library Preparation guide from Illumina, Inc. was used for library denaturation, dilution, and loading onto the Illumina MiSeq. The Illumina MiSeq Reagent Kit V2 (300-cycle) was used (Illumina, Inc.). The pooled DNA library and PhiX Control were denatured and then diluted to 4 pM. A 15% PhiX Control spike was used for quality control during sequencing. Forward and reverse read lengths were 151 base pairs each. The Q-score from the MiSeq runs were Q30 > 75%, indicating a base call accuracy of ≥ 99.9% for >75% of reads. The numbers of clusters passing filter were consistently >70%.

### 2.5 Bioinformatics and Data Analysis

Fastq files were demultiplexed by the MiSeq software using their sample-specific indices, and the resulting reads were processed in QIIME 2 (version 2019.10) running on VirtualBox (version 6.1.10). For quality control, raw reads underwent filtering, denoising, pairwise merging, and chimera removal via the DADA2 algorithm, which employs advanced denoising to resolve true amplicon sequence variants (ASVs) and minimize sequencing errors. Taxonomic assignment was then carried out against the Greengenes 13_8 reference database, and any sequences annotated as mitochondrial or chloroplast in origin were discarded. To standardize sampling depth for downstream diversity metrics, each sample’s read count was rarefied to 12,000 sequences. Phylogenetic analyses were facilitated by the q2-phylogeny plugin’s align-to-tree-mafft-fasttree workflow. Finally, alpha diversity was quantified for each sample by calculating observed Operational Taxonomic Unit (OUT) richness, Shannon diversity index (Shannon Entropy), Pielou’s evenness, Faith’s phylogenetic diversity, and Good’s coverage to evaluate sequencing completeness and community structure.

Data analyses of rhizobacterial DNA were conducted in R! All graphs were produced using the ggplot2 package. Chloroplast and mitochondrial DNA were manually removed from the January 2023-2025 and June 2023-2025 datasets (<6% of all DNA sequenced). In the January 2023-2025 experiments, alpha rarefaction plateau for Shannon entropy (8 bits) and Operational Taxonomic Unit (OTU; >700 counts) were achieved at sequencing depths of 1,000 and 4,000 reads, respectively. Alpha rarefaction plateau for Shannon entropy (8 bits) and OTU counts (>700) were achieved at sequencing depths of 500 and 4,000 reads in the June 2023-2025 experiments. PHC-degrading bacteria and genes were identified through the National Center for Biotechnology (NCBI) Gene database (Figures 10 - 11).

## 3.0 Results

### 3.1 Soil PHC Levels

CCME Fraction (F) 2-4 levels in the 25,700 mg/kg TPH soil were significantly higher than maximum recommended guidelines provided at the federal and provincial levels for coarse-grained industrial soil (Table 2: CCME 2008; OME 2011). The <120 mg/kg TPH soil contained below-threshold PHC fraction levels (Table 2). The 12,600 mg/kg TPH soil contained four-folds higher F2 hydrocarbons than federal and provincial guidelines.

**Table 2.** Petroleum hydrocarbon (PHC) fraction concentrations in the <120 mg/kg TPH (*n*=9), 25,700 mg/kg TPH (*n*=9) and 12,600 mg/kg TPH (*n*=6) soils from the contaminated site located in northern Ontario. Mean concentrations and standard deviation are provided in brackets for the site soils. PHC concentrations were identified by CCME Fractions and were compared against federal (CCME) and provincial PHC guidelines for coarse-grained soil collected from an industrial site.

|  | Fraction 1<sup>1</sup><br>(mg/kg) | Fraction 2<sup>2</sup><br>(mg/kg) | Fraction 3<sup>3</sup><br>(mg/kg) | Fraction 4<sup>4</sup><br>(mg/kg) | TPH (mg/kg) |
| --- | --- | --- | --- | --- | --- |
| <120 mg/kg TPH<br>(Background) | <10 | <10 | 27 (4.2) | 32 (5.0) | 60 (6.2) |
| 25,700 mg/kg TPH<br>(Contaminated) | <10 | 10,500<br>(683) | 11,500<br>(707) | 3,600 (207) | 25,700<br>(1,580) |
| 12,600 mg/kg TPH<br>(Diluted) | <10 | 4,070<br>(946) | 6,410<br>(592) | 2,130 (130) | 12,600<br>(1,130) |
| CCME Guidelines<br>(CCME, 2008) | 320 | 260 | 1,700 | 3,300 | 5,580 |
| Provincial<br>Guidelines (OME,<br>2011) | 55 | 230 | 1,700 | 3,300 | 5,285 |

<sup>1</sup>C6-C10 effective carbon number (ECN)
<sup>2</sup>C10-C16 ECN
<sup>3</sup>C16-C32 ECN
<sup>4</sup>C32+ ECN

### 3.5 Alpha Diversity

A three-way Kruskal-Wallis Test on Shannon entropy and operational taxonomic unit (OTU) richness revealed that neither plant species, PHC contamination level (soil type), nor treatment - nor any of their two- or three-way interactions - had a statistically significant effect on rhizobacterial alpha (within-sample) diversity in either the January 2023 - 2025 or June 2023 - 2025 experiments (Figure S8; all *p* > 0.05). In the January 2023 – 2025 experiments, <120 mg/kg TPH soil treatments supported a mean Shannon entropy of 8.7 ± 0.1 (mean and SD) bits versus 8.2 ± 0.2 bits in the 25,700 mg/kg TPH soils, indicating high bacterial evenness within the soils for all treatments. A moderate rank-biserial effect size (*r* = 0.39) was observed, indicating reduced evenness under PHC contamination, despite statistical insignificance (Mann-Whitney *U* Test: *p* = 0.09). In the June experiments, <120 mg/kg TPH soil treatments averaged 8.5 ± 0.2 bits compared to 8.8 ± 0.2 bits in the 12,600 mg/kg TPH soils, with non-significant differences (*p* = 0.21) and a small negative effect size (*r* = - 0.28).

OTU richness likewise exhibited minimal responses to soil contamination, treatment, and plant species (Figure S9). In the January 2023 - 2025 experiments, mean OTU counts were 804 ± 73 in the <120 mg/kg TPH soil and 721 ± 81 in the 25,700 mg/kg TPH soil. During the June 2023 – 2025 experiments, OTU richness averaged 787 ± 64 and 870 ± 70 in <120 mg/kg TPH and 12,600 mg/kg TPH soil treatments, respectively, indicating high taxonomic richness across soils and treatments. Treatment effects were similarly negligible in both the January (*p* = 0.51, *r* = - 0.15) and June experiments (*p* = 0.35, *r* = 0.26). Plant species influenced richness only in the January experiments, with *P. mariana* supporting significantly higher OTU counts than *A. gerardii* (*p* = 0.03, *r* = - 0.84), with a large effect size between the two species (Figure S9B).

### 3.6 Beta Diversity

A permutational multivariate analysis of variance (PERMANOVA, *n* = 999) based on Bray-Curtis Dissimilarity across the factors of soil type, plant species, and treatment revealed significantly variable beta (between-sample) diversity in the January 2023 – 2025 experiments (*F* = 3.28, *p* = 0.001, Pearson’s correlation *r* = 0.49). Pair-wise PERMANOVA tests with Bonferroni-adjusted *p*-values showed that soil type significantly influenced beta diversity (*p* = 0.001), with moderate effect size (*r* = 0.35) but treatment and plant species did not impact community diversity (*p* > 0.05 for both). Bray-Curtis Dissimilarity (0.80 – 1.0) and the two primary hierarchical dendrogram clusters indicate soil PHC contamination as the most significant factor for soil microbial community structure while plant species (0.50 – 0. 75) and treatment (0.40 - 0.75), exert non-significant secondary and tertiary effects on community composition, respectively (Figure 2).

**Figure 2.**
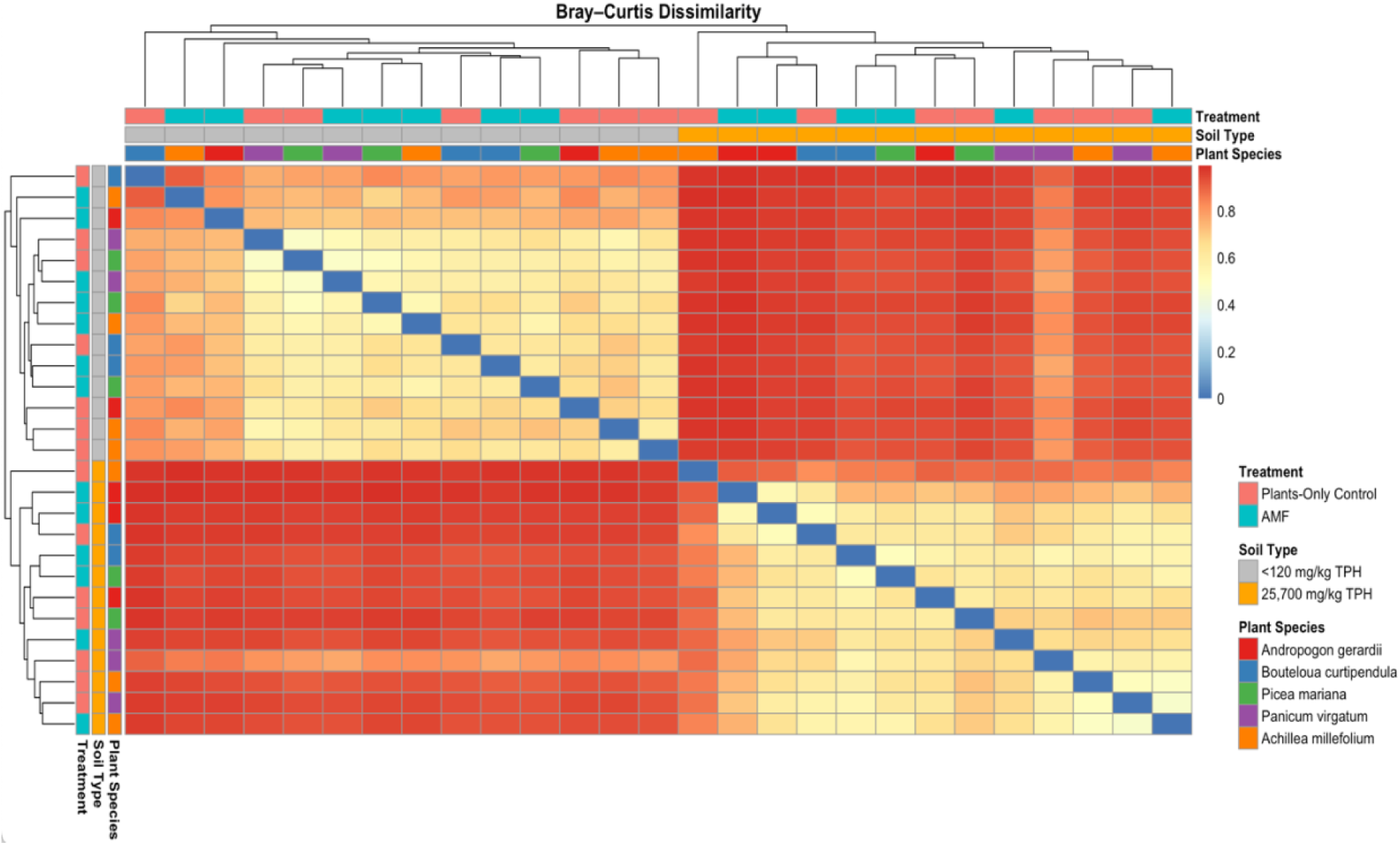
Heat map and dendrogram hierarchical cluster analysis of soil bacterial communities based on Bray-Curtis Dissimilarity. Bacterial communities were sequenced in <120 mg/kg total petroleum hydrocarbon (TPH) and 25,700 mg/kg TPH soils during the January 2023-2025 phytoremediation experiments. Two soil treatments (Plants-Only Control and AMF) were used in conjunction with five plant species (*Andropogon gerardii*, *Bouteloua curtipendula*, *Picea mariana*, *Panicum virgatum*, and *Achillea millefolium*).

A PERMANOVA (*n* = 999) of Bray-Curtis Dissimilarity applied to the June 2023 – 2025 data revealed no significant variation in beta diversity across plant species, soil type, and treatment (*F* = 1.91, *p* = 0.20, *r* = 0.20). However, pair-wise PERMANOVA with Bonferroni-adjusted *p*-values showed that soil type significantly influenced beta diversity (*p* = 0.014), with small effect size (*r* = 0.07). Treatment and plant species did not impact community diversity (*p* > 0.05 for both). Bray-Curtis Dissimilarity (0.50 – 1.0) and the two primary hierarchical clusters indicate soil PHC contamination as the most significant factor for soil microbial community structure in the June experiments (Figure 3). Soil treatment (0.45– 0.75) and plant species (0.38 – 0.70) exert secondary and tertiary effects on community composition.

**Figure 3.**
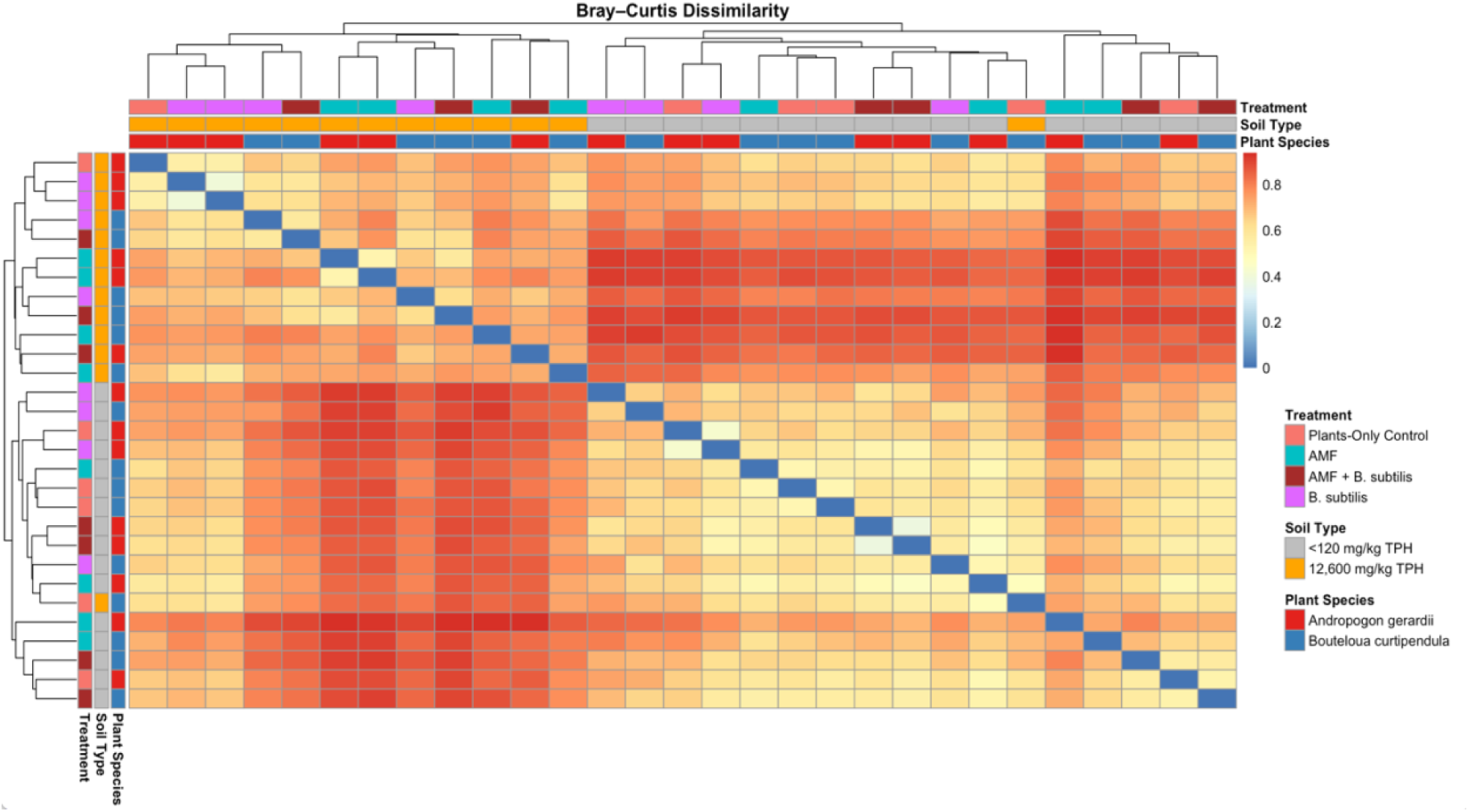
Heat map and dendrogram hierarchical cluster analysis of soil bacterial communities based on Bray-Curtis Dissimilarity. Bacterial communities were sequenced in <120 mg/kg total petroleum hydrocarbon (TPH) and 12,600 mg/kg TPH soils during the June 2023-2025 phytoremediation experiments. Four soil treatments (Plants-Only Control, AMF, *Bacillus subtilis*, and both AMF and *B. subtilis*) were used in conjunction with two plant species (*Andropogon gerardii* and *Bouteloua curtipendula*).

Principal coordinates analysis (PCoA) of Bray–Curtis dissimilarities revealed that the first axis (PCoA1) explained 36.6% of the total community variance while the second axis (PCoA2) accounted for 6.2% of the variance in the January 2023–2025 experiments, for a total contribution of 42.8% (Figure 4A). There is clear visual separation of samples based on soil type when accounting for both rare and abundant taxa. Correspondingly, principal component analysis (PCA) of central-log ratio (CLR-) normalized OTU abundances recapitulated this pattern while only accounting for the most abundant taxa: PC1 captured 10.6% of the variance, and PC2 accounted for 7.5% of the variance, for a total contribution of 18.1% (Figure 4B). Neither treatment nor plant species exhibited consistent segregation along either PCoA ordination axis (Figures S10 – S11), demonstrating that soil type is the primary factor structuring rhizobacterial community composition.

**Figure 4.**
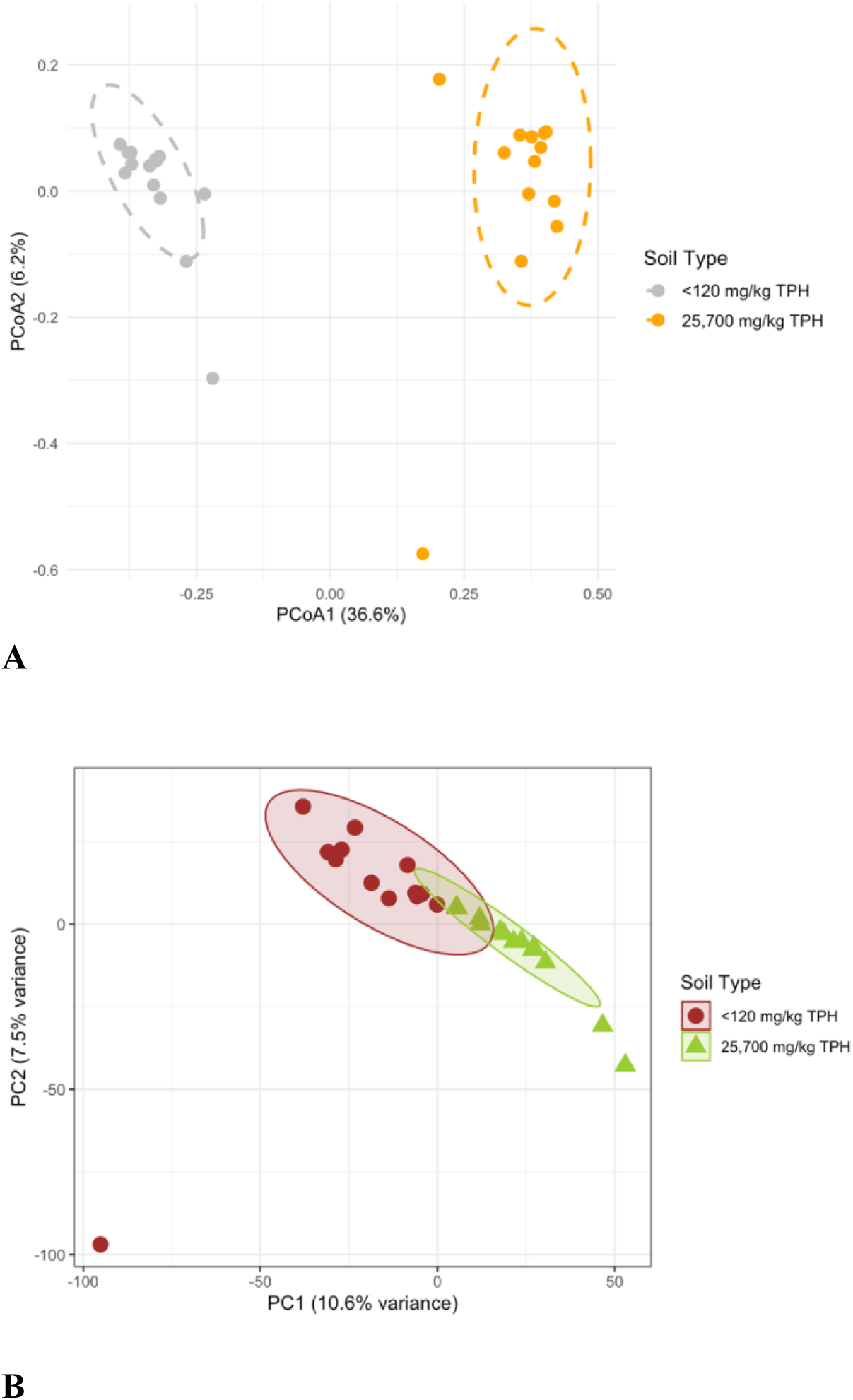
Visualization of rhizobacterial community separation based on soil type during the January 2023-2025 experiments. **A.** Principal coordinates of analysis based on soil type (<120 mg/kg TPH and 25,700 mg/kg TPH). **B.** Central-log ratio (CLR) – normalized principal component analysis based on soil type (<120 mg/kg TPH and 25,700 mg/kg TPH).

A PCoA applied to the June 2023-2025 experiments revealed that the PCoA1 explained 24.8% of the total community variance while PCoA2 accounted for 7.0 % of the variance, indicating that 31.8% of the data can be accounted for by these two ordinal axes (Figure 5A). Samples clustered distinctly by soil type, reflecting variation in both rare and abundant taxa. PCA of CLR–normalized OTU abundances—emphasizing the most abundant taxa—demonstrates more modest ordination power (PC1 = 8.9%, PC2 = 6.7% of total variance; Figure 5B), representing a total contribution of 15.6%, with no consistent clustering by treatment or plant species (Figures S12 - S13).

**Figure 5.**
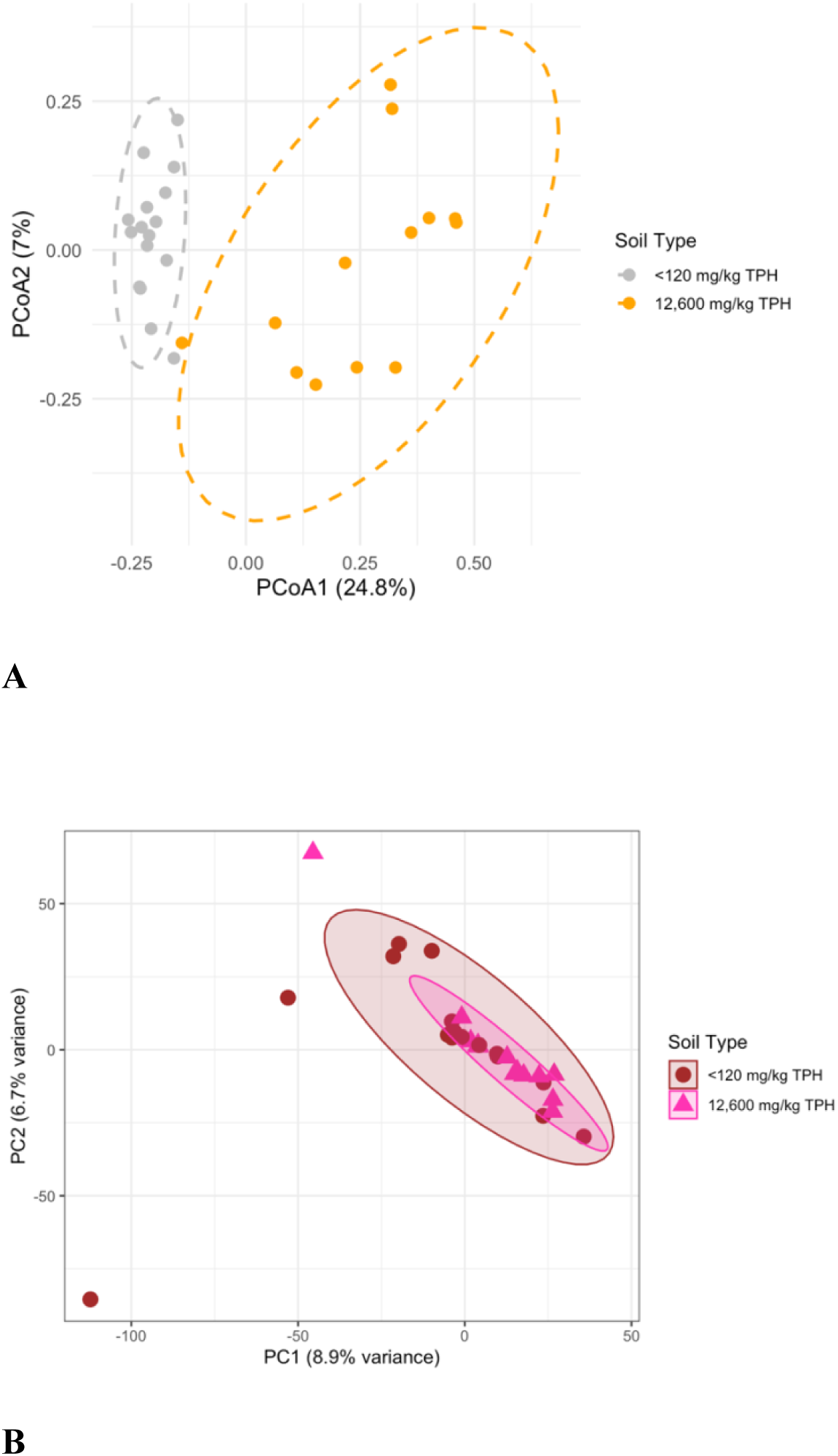
Visualization of rhizobacterial community separation based on soil type during the June 2023-2025 experiments. **A.** Principal coordinates of analysis based on soil type (<120 mg/kg TPH and 12,600 mg/kg TPH). **B.** Central-log ratio (CLR) – normalized principal component analysis based on soil type (<120 mg/kg TPH and 12,600 mg/kg TPH).

### 3.7 Bacterial Abundance in Soils

Proteobacteria was the predominant phylum in the 25,700 mg/kg TPH soil, with a total abundance of 330,000 bacteria (37.9% of all bacterial counts), including the hydrocarbon-degrading genera, *KCM-B-112* (22,100 sequences; 2.54%) and *Sphingomonas* (16,500; 1.89%) (Figure 6). Actinobacteriota ranked second most abundant at 157,000 sequences (18.0%), followed by Acidobacteriota (117,000; 13.4%) and Chloroflexi (66,000; 7.60%). Overall, the 25,700 mg/kg TPH soil exhibited 872,000 total bacteria compared with 657,000 in the <120 mg/kg TPH soil, representing a 32.8% greater bacterial load (Figures 6 – 7; Mann-Whitney *U* test: *p* = 0.04, rank biserial *r* = 0.47). The AMF treatment increased absolute bacterial abundance by 24.0%, from 818,000 to 710,000 bacterial counts in the Plants-Only Control. Finally, the native biosurfactant producer *Sporosarcina*, (phylum: Firmicutes), was significantly enriched in the 25,700 mg/kg TPH soil (9,920; 1.14%) versus the <120 mg/kg TPH soil (251; 0.03%).

**Figure 6.**
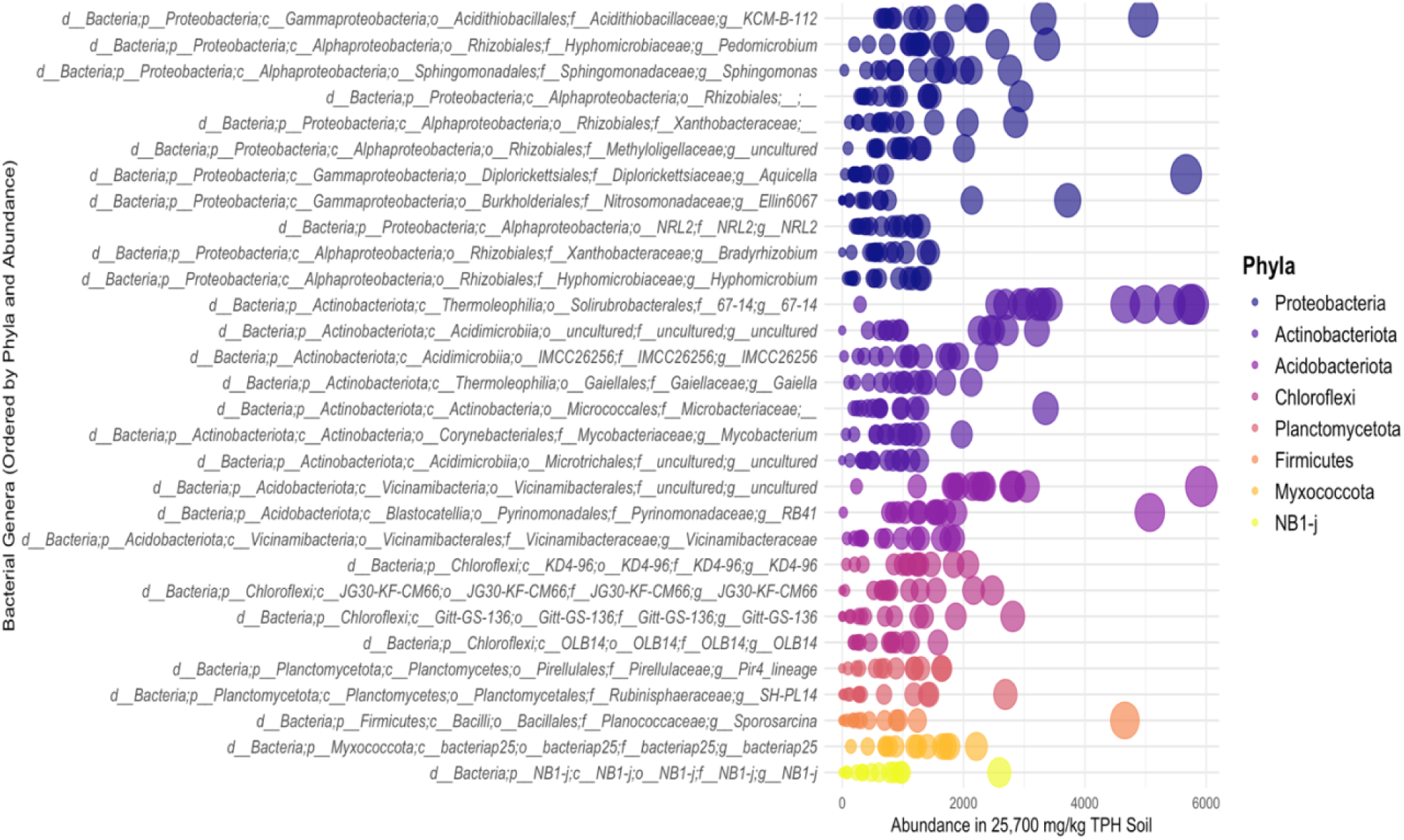
Bubble chart of the absolute abundance of the thirty most abundant bacteria found in the 25,700 mg/kg contaminated TPH soil during the January 2023-2025 experiments, representing 54.1% of all bacteria in the soil. Each bubble represents a soil sample. The y-axis indicates taxonomic classification from domain (d) to genera (g). Genera are colour-coded by phyla classification in the legend. The legend presents phyla by absolute abundance in descending order.

Within the <120 mg/kg TPH soil, Actinobacteriota was the dominant phylum, contributing to a bacterial count of 212,000 (32.3% of all bacteria). The hydrocarbon-degrading genus, *KCM-B-112*, was strongly enriched in the 25,700 mg/kg TPH soils, expanding 103-fold (2.54% vs. 0.025%), whereas *Sphingomonas* showed only a modest 9.2% increase (1.89% vs. 1.73%). In the 25,700 mg/kg TPH soil, Proteobacteria ranked first in absolute abundance (331,000, 37.9%), followed by Actinobacteriota (157,000; 18.0%) and Acidobacteriota (117,000; 13.4%). In the <120 mg/kg TPH soil, AMF inoculation boosted absolute bacterial abundance by 58.0% - from 402,000 to 254,000 relative to the Plants-Only Control. Finally, the nitrogen-fixing phylum, NB1-j, and the predatory phylum, Myxococcota, were 22.1-fold and 1.84-fold more abundant, respectively, in the 25,700 mg/kg TPH soils compared to the <120 mg/kg TPH soils (Figures 6 - 7).

**Figure 7.**
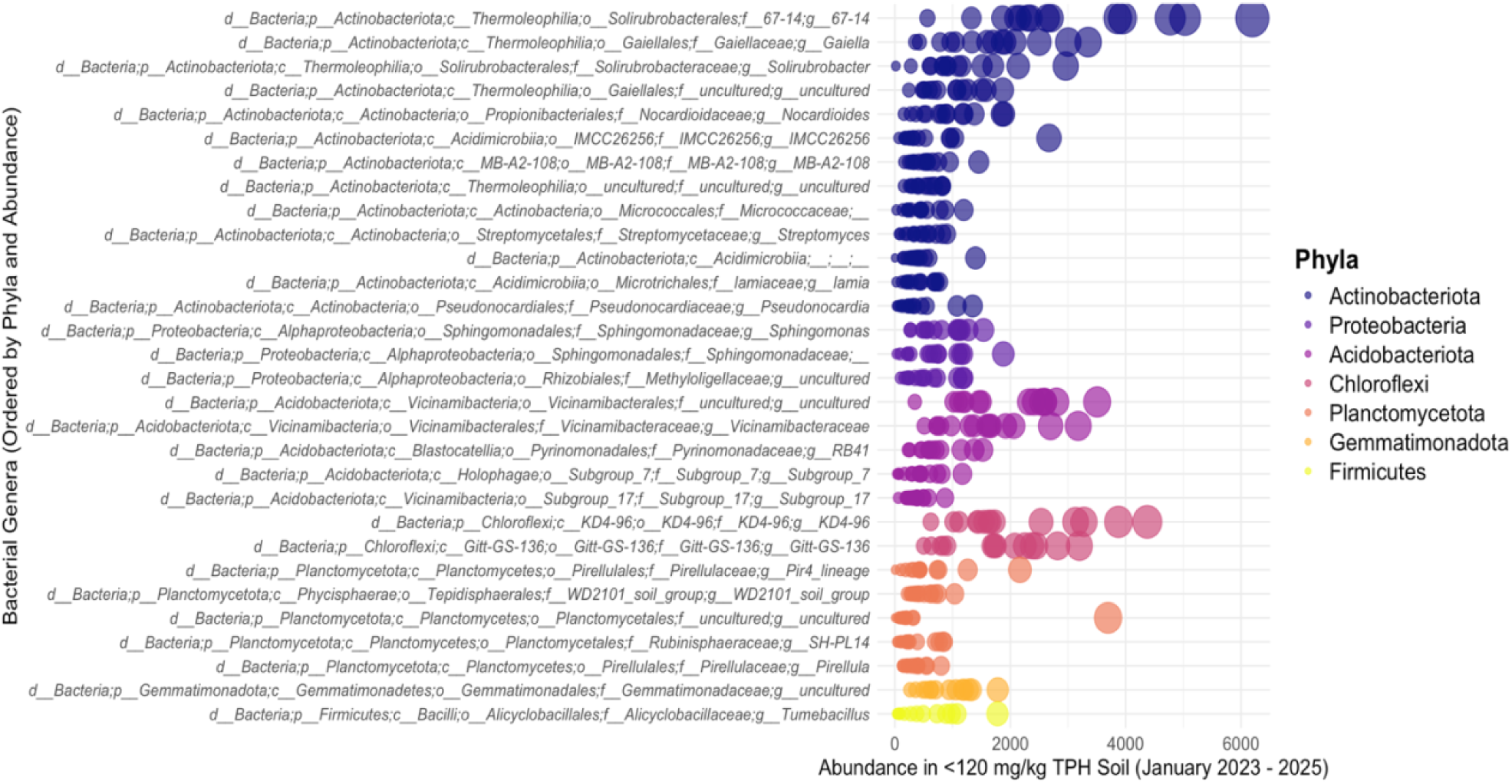
Bubble chart of the absolute abundance of the thirty most abundant bacteria found in the <120 mg/kg TPH background soil during the January 2023-2025 experiments, representing 61.2% of all bacteria in the soil. The y-axis indicates taxonomic classification from domain (d) to genera (g). Genera are colour-coded by phyla classification in the legend. The legend presents phyla by absolute abundance in descending order.

In the 12,600 mg/kg TPH soil, Proteobacteria remained the dominant phylum (Figure 8), accounting to a bacterial count of 246,000 (28.2% of all bacteria). Actinobacteriota was the second most abundant at 153,000 reads (17.7%), followed by Acidobacteriota with 148,000 (17.0%) and Chloroflexi with 86,900 (10.1%). Key hydrocarbon-degrading genera comprised *Sphingomonas* (15,100 reads; 1.75%), *Mycobacterium* (8,760; 1.01%), *KCM-B-112* 7 (7,250; 0.84%), and *Rhodococcus* (521; 0.06%). Compared to the 25,700 mg/kg soil, Acidobacteriota and Chloroflexi were 27.1% and 31.3% more abundant, whereas *KCM-B-112* was three-fold more enriched under the 25,700 mg/kg TPH soils. *Mycobacterium* and *Sphingomonas* were 28.0% and 9.4% more enriched in the 25,700 mg/kg TPH soil, respectively. In contrast, *Rhodococcus* was three-fold more abundant in the 12,600 mg/kg TPH soil. There was no significant difference in total PHC-degrader abundance between soils (Mann–Whitney *U* test, *p* = 0.52).

**Figure 8.**
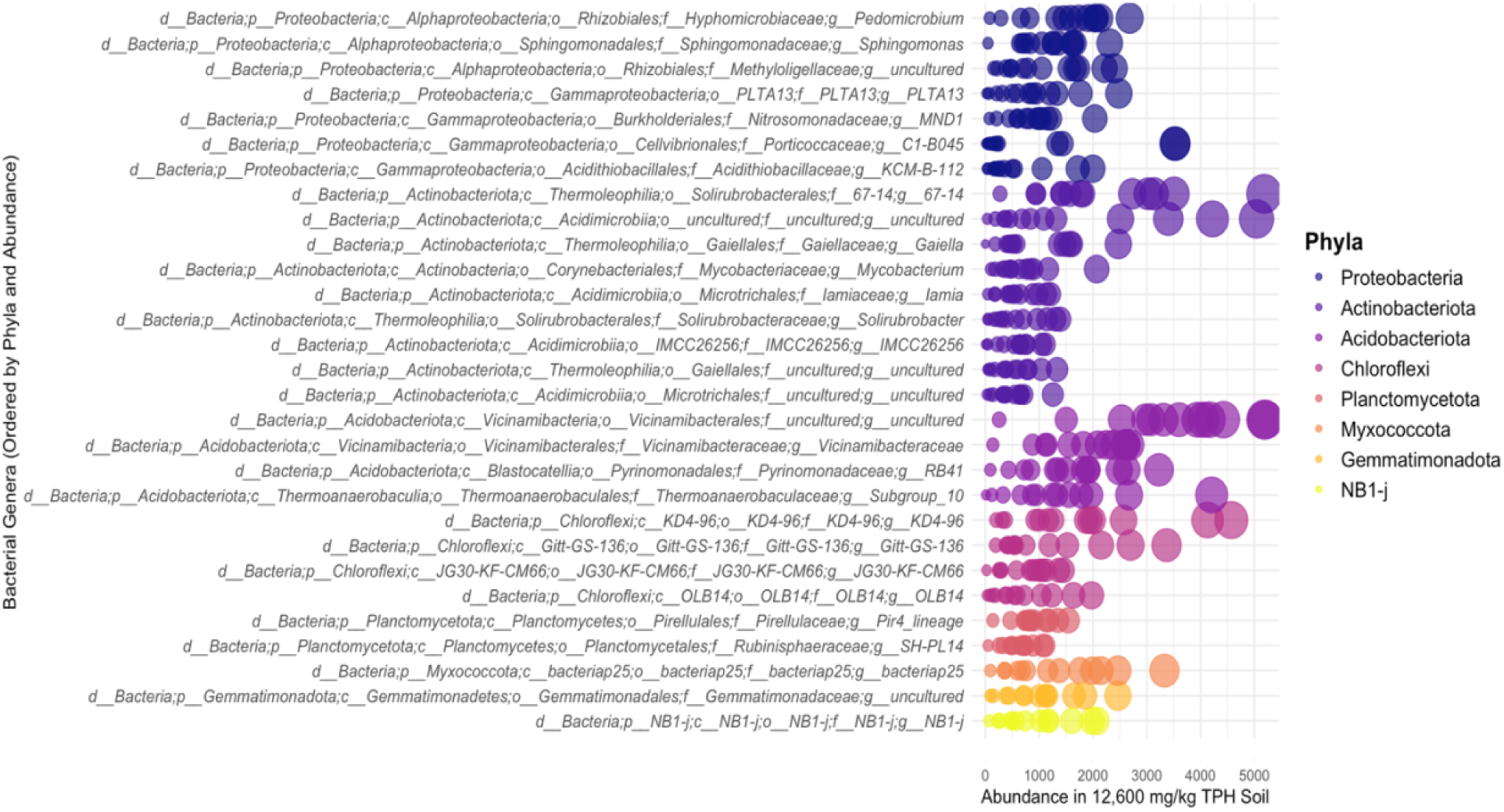
Bubble chart of the absolute abundance of the thirty most abundant bacteria found in the 12,600 mg/kg TPH diluted soil during the June 2023-2025 experiments, representing 48.9% of all bacteria in the soil. Each bubble represents a soil sample. The y-axis indicates taxonomic classification from domain (d) to genera (g). Genera are colour-coded by phyla classification in the legend. The legend presents phyla by absolute abundance in descending order.

Neither *B. subtilis* nor its broader genus or phylum reached notable abundance in the diluted soil despite inoculation (Figure 8), suggesting competitive exclusion (3,690 bacterial count; 0.43%). The *B. subtilis* treatment contained the highest absolute abundance of the species (297,000; 34.1% relative abundance across treatments), followed by AMF (234,000; 26.8%), AMF + *B. subtilis* (182,000; 20.9%), and the Plants-Only Control (151,000; 17.3%). No significant differences in *B. subtilis* abundance were detected across treatments (Kruskal-Wallis *H* = 4.092, *p* = 0.252, *df* = 3).

Bacterial abundance was not significantly different between the 12,600 mg/kg TPH soil and <120 mg/kg TPH soil for the June 2023-2025 experiments (864,000 vs 934,000 total bacterial counts; Mann-Whitney *U* test: *p* = 0.42). In the 12,600 mg/kg TPH soil, Proteobacteria was the most abundant phylum (246,000; 28.5% of all bacteria), followed by Actinobacteriota (153,000; 17.7%), Acidobacteriota (148,000; 17.2%), and Chloroflexi (86,900; 10.1%) (Figure 9). *KCM-B-112* was 50-fold more enriched under the 12,600 mg/kg TPH soil (7,250 vs 146 bacterial counts). *Mycobacterium* was three-fold more enriched in the 12,600 mg/kg TPH soil. (8,760 vs. 2,820). *Rhodococcus* abundance did not differ between the <120 mg/kg TPH and 12,600 mg/kg TPH soils (15,600 vs. 15,100). *B. subtilis* was detected in all soil-inoculation treatments in the <120 mg/kg TPH soil, with the highest *Bacillus* genera load under the Plants-Only Control (6,740; 0.72%), followed by the AMF + *B. subtilis* treatment (2,840; 0.30%), the *B. subtilis* treatment (2,500; 0.27%), and the AMF treatment (2,220; 0.24%).

**Figure 9.**
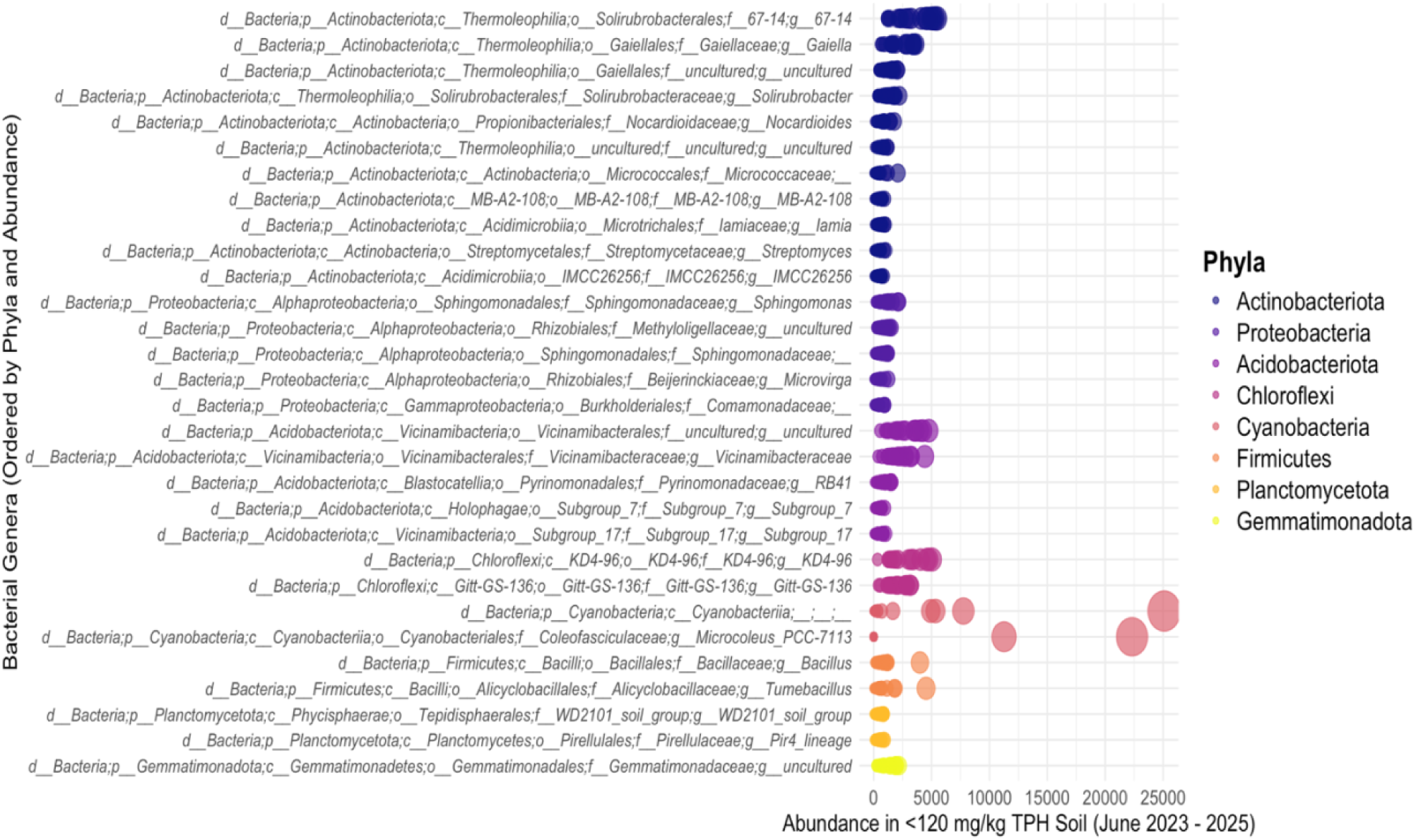
Bubble chart of the absolute abundance of the thirty most abundant bacteria found in the <120 mg/kg TPH background soil during the June 2023 - 2025 experiments, representing 58.9% of all bacteria in the soil. Each bubble represents a soil sample. The y-axis indicates taxonomic classification from domain (d) to genera (g). Genera are colour-coded by phyla classification in the legend. The legend presents phyla by absolute abundance in descending order.

### 3.8 Bacterial Functional Analyses

Thirteen PHC-degrading bacterial phyla were identified in the January 2023-2025 and June 2023-2025 experiments through NCBI Genes (Figures 10 - 11). Three alkane-degrading genes were identified: *alkB* (encoding alkane monooxygenase), *CYP153* (cytochrome P450 alkane hydroxylase), and *assA* (alkylsuccinate synthase). *Alkb* and *CYP153* are responsible for aerobic degradation of aliphatic PHCs while *assA* is involved in anaerobic degradation. *Alkb* was found in all phyla except Bacteroidota, Acidobacteriota, and Desulfobacterota. Alkylsuccinate synthase is only found in Desulfobacterota. Thirteen aromatic-degrading genes were identified (Figures 10 - 11).

**Figure 10.**
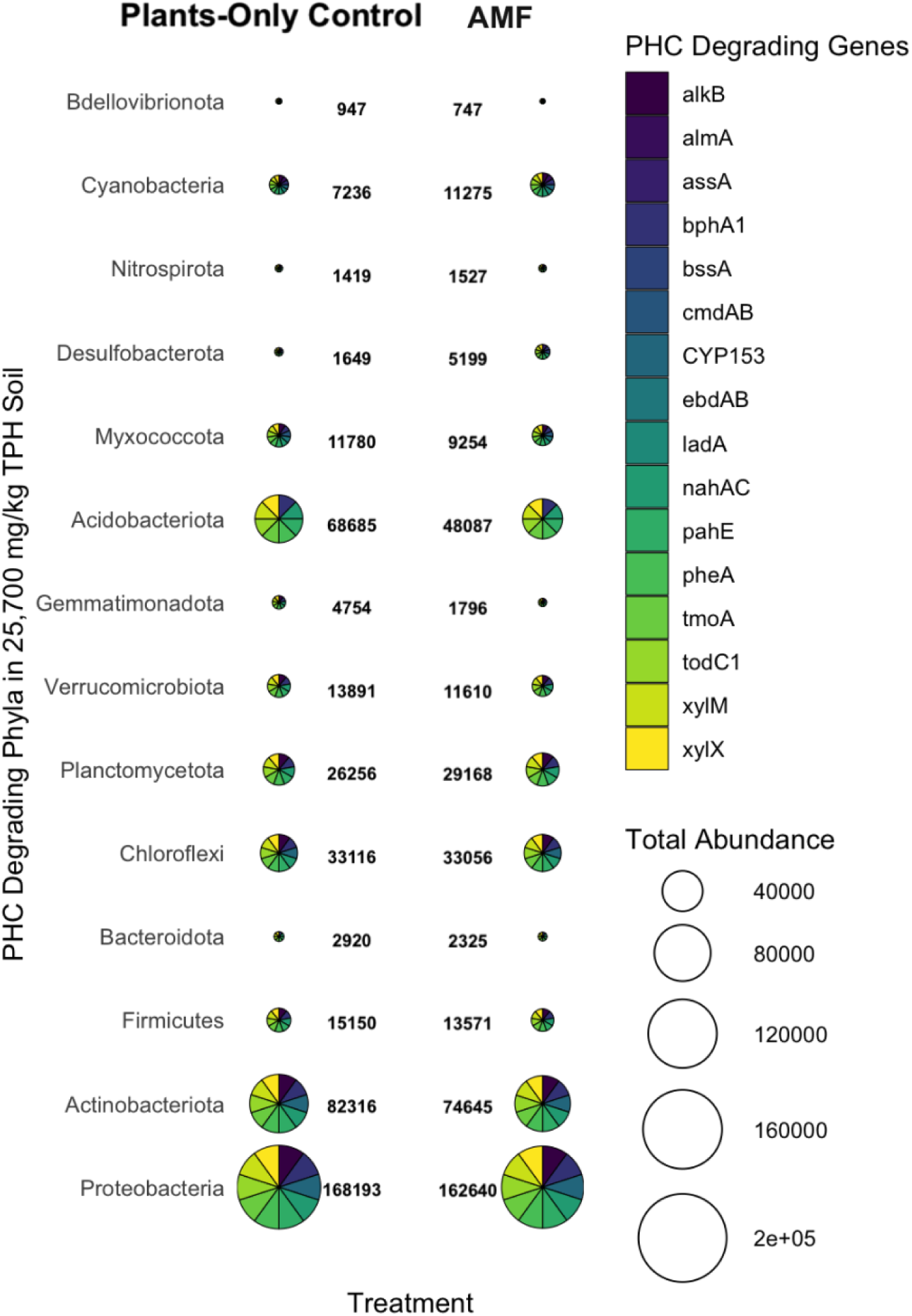
Distribution and total abundance counts of PHC-degrading bacterial phyla in the 25,700 mg/kg TPH soil from the January 2023 – 2025 experiments. Scatterpie bubble charts illustrate phylum-level bacterial communities under two treatments: Plants-Only Control and AMF inoculation across five phytoremediation plant species (*Andropogon gerardii*, *Bouteloua curtipendula*, *Panicum virgatum*, *Achillea millefolium*, and *Picea mariana*). Bubble size corresponds to total bacterial abundance within each phylum (numerical values displayed adjacent to bubbles). Pie segments represent the distribution of PHC-degrading genes identified within each bacterial phylum based on functional gene annotation.

**Figure 11.**
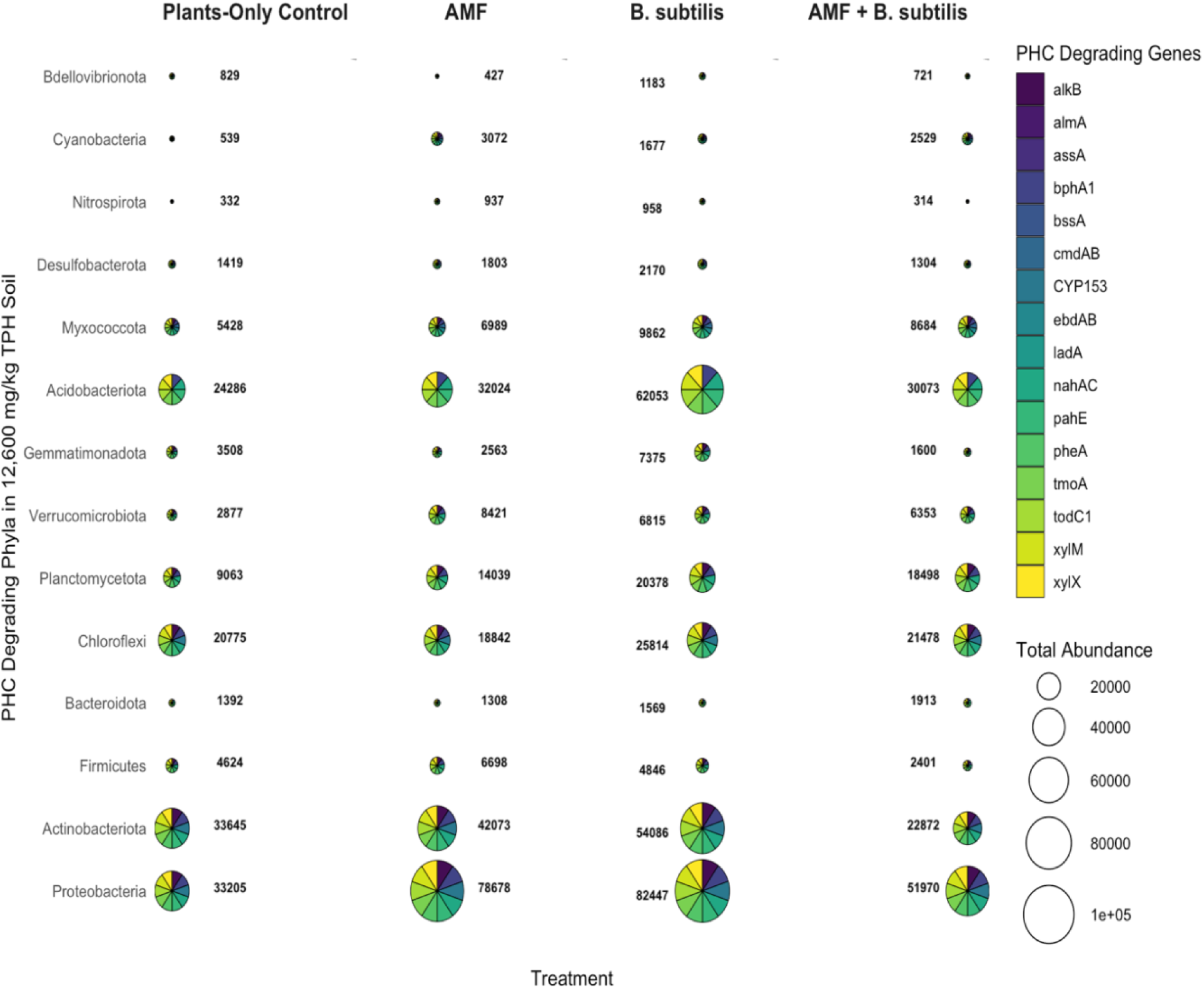
Distribution and total abundance counts of PHC-degrading bacterial phyla in the 12,600 mg/kg TPH soil from the June 2023 – 2025 experiments. Scatterpie bubble charts illustrate phylum-level bacterial communities under four treatments: Plants-Only Control, AMF, *Bacillus subtilis* (*B. subtilis*), and AMF + *B. subtilis* soil inoculation across two phytoremediation plant species (*Andropogon gerardii* and *Bouteloua curtipendula*). Bubble size corresponds to total bacterial abundance within each phylum (numerical values displayed adjacent to bubbles). Pie segments represent the distribution of PHC-degrading genes identified within each bacterial phylum based on functional gene annotation.

Ten of the aromatic-degrading genes indicated in Figures 10 - 11, excluding *bssA* (benzylsuccinate synthase), *ebdAB* (ethylbenzene dehydrogenase), and *cmdAB* (p-cymene dehydrogenase), encode dioxygenases and monooxygenases that partake in aerobic degradation pathways. The following aromatic-degradation genes were found in all phyla: *nahAC*, *bphA1*, *pheA*, *xylM*, *xylX*, *todC1*, *tmoA*, *pahE*. Myxococotta is the only phylum that possesses all three of the anaerobic aromatic-hydrocarbon degrading genes, while Desulfobacterota possesses the *bssA* gene. All soils contained significantly higher aerobic relative to anaerobic PHC-degrading genes (Mann-Whitney U tests: *p* < 0.05), with very high effect size for all soils, treatments, and plant species (rank biserial *r* = 1.0 in all comparisons).

There was no significant difference observed between total abundance of PHC-degrading bacteria in the 25,700 mg/kg TPH and 12,600 mg/kg TPH soils (Mann–Whitney *U* test, *p* = 0.90). The total abundance of PHC-degrading bacterial phyla was significantly higher in the 25,700 mg/kg TPH contaminated soil compared to the <120 mg/kg TPH background soil in the January 2023 – 2025 experiments (*p* = 0.04), whereas the 12,600 mg/kg TPH soil did not differ significantly from the <120 mg/kg TPH soil (*p* = 0.58) in the June 2023 - 2025 experiments (Figure S14). In the 25,700 mg/kg TPH soil, Proteobacteria remained the dominant PHC-degrading phylum under both treatments, followed by Actinobacteriota, and Acidobacteriota (Figure 10). Mann–Whitney *U* tests revealed no statistically significant difference in mean abundance between the Plants-Only Control and AMF treatments (all *p* > 0.05) for all PHC-degrading phyla in the soil samples. Likewise, there was no significant difference in aerobic or anaerobic PHC-degrading gene abundance between the two treatments in the 25,700 mg/kg TPH soil (both *p* > 0.05).

Kruskal–Wallis and Mann-Whitney *U* tests were applied to compare the total abundance of each PHC-degrading bacterial phylum across the four treatments in the 12,600 mg/kg TPH soil. No phylum exhibited a statistically significant difference between treatments for all test (all *p* > 0.05; Figure 11). Neither the AMF nor the *B. subtilis* treatment showed a statistically significant increase in Proteobacteria abundance compared to the Plants-Only Control or to the AMF + *B. subtilis* treatment (Mann-Whitney *U* tests: all *p* > 0.05), despite a 2.3 – 2.5 fold increases relative to the Plants-Only Control and a 51% and 59% increase relative to the AMF + *B. subtilis* treatment. There was no significant difference in aerobic or anaerobic gene abundance between the different treatments for both Kruskal–Wallis and Mann-Whitney *U* tests (all *p* > 0.05).

## 4.0 Discussion

### 4.1 PHC Contamination Drives Rhizobacterial Biodiversity

This study revealed that soil PHC contamination is the primary driver of rhizobacterial community composition during rhizodegradation (Figures 2 - 5). The magnitude of PHC contamination also significantly influences rhizobacterial composition in soil. The fully contaminated site soil (25,700 mg/kg TPH) resulted in more distinct microbial communities relative to the background soil (<120 mg/kg TPH) in the January 2023 – 2025 experiments (Figure 4), compared to the diluted site soil (12,600 mg/kg TPH) and background soil (<120 mg/kg TPH) from the June 2023 – 2025 experiments (Figure 5). Several studies examining the microbiome have shown that PHC contamination is a dominant environmental factor structuring microbial community composition in various environmental matrices (Paissé et al., 2008; Jia et al., 2023), with higher soil PHC contamination more significantly varying community structure (Abed et al. 2015). In chronically oil-polluted sediments, PHC contamination level has been shown to influence bacterial community structure more than other environmental factors (Paissé et al., 2008). Jia et al. (2023) demonstrated that PHCs alone accounted for 45.8% of the microbial community diversity – higher than soil physicochemical properties (26.4%) and soil lithology (0%).

Phytoremediation plant species or soil inoculation treatment (Plants-Only Control, AMF, *B. subtilis*, AMF + *B. subtilis*) did not significantly influence rhizobacterial community composition through Shannon entropy, OTU, or Bray-Curtis Dissimilarity (Figures 2 - 3; S8 - S9). Alpha diversity was high in all soils (Figures S8 - S9), which may be reflective of single plant inoculation enhancing bacterial biomass of all soils (Schlatter et al., 2015). Schlatter et al. (2015) demonstrated an inverse relationship between plant species richness and rhizobacterial diversity. Both plant and rhizobacterial growth is limited by soil carbon availability, with more variable carbon sources and quantity diversifying bacterial diversity (Grayston et al., 1998). Nutrient analyses revealed that the non-planted <120 mg/kg TPH background soil utilized in this study are carbon-deficient (1.49% organic C) while the 25,700 mg/kg TPH contaminated soil is not (7.79% organic C; Table S2), suggesting that PHC contamination may have enhanced rather than constrained microbial diversity. Significance testing between the 25,700 mg/kg TPH soil and the <120 mg/kg TPH soil from the January 2023 – 2025 experiments confirmed that total abundance of total bacterial abundance was significantly enhanced under PHC contamination (Mann-Whitney *U* test: *p* = 0.04).

Plant species have been shown to significantly alter rhizobacterial community composition based on phylogenetic distance of plant species and plant root exudate composition (Marschner et al., 2004; Garbeva et al., 2008; Lei et al., 2018). Other studies have shown that plants support similar microbial communities in various uncontaminated soils (Grayston et al., 1998; Miethling et al., 2000), which may be attributed to similar carbon availability in the soils. Dagher et al. (2019) showed that PHC concentration levels may have stronger influence on bacterial community composition than plant species or treatment (Proteobacteria consortia bioaugmentation) during remediation of PHC-contaminated sediments. This suggests that PHC contamination is the most significant driver of microbial community composition during rhizodegradation of PHCs in both soil and sediment matrices.

### 4.2 Bacterial Abundance in Soils

The most abundant bacterial phyla across the site soils (Proteobacteria, Actinobacteriota, and Acidobacteriota) are likely reflective of indigenous rhizobacteria that contribute to soil nutrient cycling and organic matter decomposition because they are widely found in soils from various environments (Figures 6 - 9; Spain et al. 2009; Lei et al., 2018). The fourth most abundant phyla in this study, Chloroflexi, is abundant in cold and moist environments, including soil collected from the boreal ecozone, due to their oligotrophic strategy facilitating carbon and nitrogen acquisition in nutrient-deficient soils during freeze-thaw cycles (Costello and Schmidt, 2006; Vuillemin et al., 2020). There were significantly higher Proteobacteria abundance in the 12,600 mg/kg TPH and 25,700 mg/kg TPH soils compared to the <120 mg/kg TPH soils (Figure 10), aligning with findings from microbiome studies performed on aged PHC-contaminated soils (Viñas et al., 2005; Shahi et al., 2016; Jia et al., 2023). Proteobacteria possess a broad suite of alkane monooxygenases (*alkB*, *CYP153*) and dioxygenases (*bphA1*, *nahAC*, *todC1*) enabling both aliphatic and aromatic hydrocarbon degradation (Figures 10 - 12). Oligotrophic hydrocarbon-degrading Proteobacteria, *KCM-B-112* and *Sphingomonas*, were significantly more abundant in the 25,700 mg/kg TPH soils than the <120 mg/kg TPH soils (Figures 6 - 7), as is typical of PHC-contaminated soils that have not been supplemented with added nutrients (Viñas et al., 2005; Zhou et al. 2016, 2022; Brzeszcz et al., 2023; Cagle et al., 2024). Viñas et al. (2005) reported that neither biosurfactant addition nor bioaugmentation altered PHC degradation kinetics as significantly as soil nutrient availability.

Garbisu et al. (2017) demonstrated that bioaugmentation is often unsuccessful in altering rhizobacterial community composition due to rapid decreases in bacterial abundance following inoculation, aligns with our observed absence of *B. subtilis* in the 12,600 mg/kg TPH soil (Figure 8). *Bacillus* was abundant in the <120 mg/kg TPH soils, indicating that soil inoculation was successful (Figures 9, S1, S3). However, the *B. subtilis* was likely outcompeted in the soil (Hazarika et al., 2019; Bolivar-Anillo et al., 2021). Hemocytometry and MPN assays confirmed that *B. subtilis* grew more proficiently with sucrose as a carbon source, compared to the PHC – hexadecane (Figure S3), aligning with studies that have confirmed preference of *B. subtilis* for carbohydrate carbon sources (Warda et al., 2016; Buffing et al., 2018). Culturing a similar strain of *B. subtilis* - also possessing a hydrophilic cell surface - in hexadecane, as opposed to sucrose, has shown minimal improvement in adherence of the bacteria to hydrocarbons (Chtioui et al., 2010; Wang et al., 2019), and would have risked further contaminating the site soils.

*KCM-B-112* was also abundant in the 12,600 mg/kg TPH soil (Figure 8). *KCM-B-112* can efficiently metabolize hydrocarbons since they possess genes encoding both alkane monooxygenases (*alkB, cyp153*) and ring-hydroxylating dioxygenases (e.g., *nahAC*, *bphA1*, *xylM*, *todC1*), whereas *B. subtilis* relies almost exclusively on biosurfactant production and cytochrome p450 monooxygenases in PHC-contaminated environments (Zylstra et al., 1997; Liu et al., 2004; Parthipan et al., 2017; Zhao et al., 2022). It is also probable that *B. subtilis* was preyed on by bacteria of the predatory phylum, Myxococotta, which was significantly more abundant in the 12,600 mg/kg TPH soils compared to the <120 mg/kg TPH soils (Figures 8 - 9). Myxococotta has been documented to prey on *B. subtilis* and induce sporulation in the species over vegetation (Müller et al., 2014; Muñoz-Dorado et al., 2016).

### 4.3 Soil Functional Profiling

Soil treatment did not influence the abundance of PHC-degrading bacteria in the soils (Figures 10 – 11). Soil microbial communities often harbor high functional redundancy, meaning multiple, taxonomically distinct taxa encode the same hydrocarbon-degrading pathways (Chen et al., 2022). When soil fungal and bacterial soil treatments are introduced, indigenous PHC-degrading bacteria with overlapping metabolic capabilities often maintain overall degrader abundance and activity, buffering against community shifts. Functional redundancy has been observed in various soils with fungal amendments, with high convergence across bioregions despite widely differing fungal communities (Talbot et al., 2014). Mycorrhizal fungal enzyme activity is correlated with soil chemistry and resource availability, with carbon, nitrogen, and phosphorous soil enrichment enhancing enzymatic (carbohydrase, phosphatase, chitinase, protease) activity for rapid carbon and nitrogen cycling through stimulation of rhizobacteria (Carrino-Kyker et al., 2022). The 25,700 mg/kg TPH site soil in this study is weakly acidic (Table S2: pH of 6.4), which may have reduced fungal hydrolytic enzyme activity and enzyme efficiency due to polyphenol complexation (Prieto-Rubio et al., 2024).

Furthermore, long-term contamination, as is the case for the 25+ years weathered contaminated site in this study, fosters the selection of robust degrader consortia adapted to high PHC levels (Jiao et al., 2019). These indigenous bacteria, typically Proteobacteria and Actinobacteriota, exhibit resistance (unchanged composition under disturbance) and resilience (rapid recovery) soil amendments, thereby dampening treatment effects on abundance. Jaio et al. (2019) reported *Bacillus* enrichment in PHC (*n*-octadecane) contaminated soil, followed by a decrease in abundance following remediation, with other indigenous populations rapidly proliferating with lower toxicity levels. The higher relative abundance of Burkholderia (Order: Burkholderiales) and *Rhodococcus* in the 12,600 mg/kg TPH soil (Figure 8), but not the 25,700 mg/kg TPH soils (Figure 6), and the prominence of the nutrient-cycling phylum, Gemmatimonadota, in the <120 mg/kg TPH and 12,600 mg/kg TPH soils (Figures 8 - 9), may be explained by lower hydrocarbon tolerance thresholds (Bayatian et al., 2025). Burkholderia are PHC-degraders that have shown tolerance to <20,000 mg/kg TPH soils (Hassan et al. 2015). *Rhodococcus* is another PHC-degrader that has shown mixed responses in contaminated soils – ranging from out-competition following bioaugmentation in 12,000 mg/kg TPH soil in some species to resilience in 20,000 – 30,000 mg/kg TPH soils in others (Płociniczak et al., 2020; Huang et al., 2022; Chen et al., 2023).

Finally, the aliphatic and aromatic hydrocarbon-degrading genes reported in this study (including *alkB*, *assA*, *nahAc*, *pheA*, *xylM*, *xylE, todC1*, *bphA1,* and *pahE*) are widely distributed via horizontal gene transfer (HGT) among bacterial communities, facilitating bacterial evolution and adaptation to contaminated environments (Figures 10 - 11; Shahi et al., 2017). HGT of these genes typically occur on mobile genetic vehicles (plasmids, transposons, and integrative conjugative elements), allowing rapid transfer of genetic materials between proximal bacterial cells (Ma et al., 2006; Sobecky et al., 2009; Phale et al., 2019). HGT helps ensure that PHC-degradative functional repertoire remains constant in soils, further stabilizing the abundance of indigenous PHC-degrading bacteria (Dombrowski et al., 2022). Several studies have reported increased HGT of *phnAc*- and *nahAc*-like genes, which encode the iron sulfur protein subunits of PAH dioxygenases, in petroleum contaminated sites (Herrick et al., 1997; Laurie and Jones, 2000; Wilson et al., 2003). An NCBI Gene search revealed that the *nahAc* gene was observed in all PHC-degrading phyla in this study (Figures 10 - 11), affirming the likelihood that HGT contributed towards stabilization of PHC-degrading genes among the soil treatments.

## 5.0 Conclusion

This study demonstrates that soil PHC contamination level is the predominant determinant of rhizobacterial community composition and functional potential during rhizodegradation. The 25+ years weathered contaminated soil (25,700 mg/kg TPH) from a boreal Canadian site exhibited more distinct microbial assemblages and higher absolute bacterial abundance compared to background (<120 mg/kg TPH) and diluted (12,600 mg/kg TPH) soils. Alpha and beta diversity metrics revealed that neither plant species nor soil inoculation treatments (AMF, *B. subtilis*, or their combination) significantly influenced microbial diversity or community structure beyond the overriding effect of PHC concentration. Taxonomic analyses identified Proteobacteria, Actinobacteriota, Acidobacteriota, and Chloroflexi as the dominant phyla across all soils, with hydrocarbon-degrading genera such as *KCM-B-112* and *Sphingomonas* enriched in contaminated soils – particularly the 25,700 mg/kg TPH site soil, reflecting their genetic capacity for aerobic alkane and aromatic hydrocarbon degradation. Functional gene profiling confirmed widespread distribution of key PHC-degrading genes - including *alkB*, *assA*, *nahAc*, *pheA*, *xylM*, *xylE, todC1*, *bphA1,* and *pahE* - suggesting the role of horizontal gene transfer and functional redundancy in stabilizing degradative potential against soil amendments. The prevalence of exogenously inoculated *B. subtilis* and negligible treatment effects on indigenous degrader abundance highlight the resilience and competitive dominance of native microbial communities adapted to chronic PHC stress. These findings underscore the critical importance of PHC contamination magnitude in shaping rhizobacterial dynamics and suggest that enhancing native microbial capacity, rather than bioaugmentation, may be a more effective strategy for bioremediation of aged PHC-contaminated soils. Future studies should investigate the long-term stability and functional resilience of indigenous bacterial communities under field conditions, the effects of PHC contamination and soil inoculants on plant health, and explore targeted amendments (e.g., nutrient supplementation) to further optimize *in situ* phytoremediation efficacy in boreal soils.

## Acknowledgements

We thank members of the Analytical Service Unit at Queen’s University, including Dr. Graham Cairns, Paula Whitley, and Mesha Thompson, for analytical assistance. We also thank Linda Eastcott, Dr. Kela Weber, Dr. Irish Koch, Dr. Juliana Ramsay, Candace Scott, Dr. Stephen Brown, and Taylor Vereecken for their guidance and/or support in conducting this research. Finally, we thank Matthew Francis, Officer Courtney Drew, and Officer Victoria Huppe for their laboratory assistance.

## Author Contributions

Conceptualization: Prama Roy, Dani Silveira, and Barbara Zeeb; Methodology: Prama Roy, Jenika Hazell, and Barbara Zeeb; Software: Prama Roy and Dani Silveira; Validation: Prama Roy and Dani Silveria; Formal analysis: Prama Roy and Dani Silveira; Investigation: Prama Roy, Jenika Hazell, and Dani Silveira; Resources: Barbara Zeeb, Data Curation: Dani Silveira and Prama Roy; Writing-Original Draft: Prama Roy; Writing - Review & Editing: Prama Roy, Dani Silveira, Jenika Hazell, and Barbara Zeeb; Visualization: Prama Roy; Supervision: Barbara Zeeb; Project administration: Prama Roy and Barbara Zeeb. Funding acquisition: Barbara Zeeb.

## Funding

This research was funded by the Natural Sciences and Engineering Research Council of Canada Collaborative Research and Development Grant to Dr. Barbara Zeeb at the Royal Military College of Canada (grant # 514935–17).

## Notes

### Competing Interest Statement

The authors have declared no competing interest.

