## Supplemental Table S1 to S14 for "Petroleum Hydrocarbon Concentration as the Primary Driver of Soil Bacterial Community Dynamics During Mycorrhizal Fungi and Rhizobacteria Assisted Phytoremediation"

**Table S1**. Particle-size distribution of the <120 mg/kg TPH, 12,600 mg/kg TPH, and 25,700 mg/kg TPH soils (ASTM 2014, 2021).

| **Particle-Size Fraction** | **Size Range (mm)** | **<120 mg/kg TPH** | **12,600 mg/kg TPH** | **25,700 mg/kg TPH** |
| --- | --- | --- | --- | --- |
| **Clay** | <0.005 | 6.2% | 3.5% | 5.5% |
| **Silt** | 0.005–0.075 | 18% | 27% | 17% |
| **Fine sand** | 0.075–0.425 | 25% | 32% | 14% |
| **Medium sand** | 0.425–2.0 | 28% | 24% | 25% |
| **Coarse sand** | 2.0–4.75 | 19% | 12% | 25% |
| **Gravel** | >4.75 | 3.3% | 1.6% | 13% |

**Table S2.** Nutrient and chemical properties analyzed in the uninoculated background and contaminated site soils used.

| **Content** | **Soil ID** | |
| --- | --- | --- |
|  | **<120 mg/kg TPH** | **25,700 mg/kg TPH** |
| **Ammonium- N (mg/kg)** | 1.98 | 0.49 |
| **Nitrate-N (mg/kg)** | 5.83 | 0.86 |
| **Nitrite-N (mg/kg)** | 0.13 | 0.06 |
| **P (mg/L)** | 2.9 | 2 |
| **K (mg/L)** | 240 | 170 |
| **Mg (mg/L)** | 300 | 350 |
| **Na (mg/L)** | 27 | 29 |
| **Ca (mg/L)** | 2,400 | 1,100 |
| **Total Carbon (%)** | 1.77 | 9.46 |
| **Organic Carbon (%)** | 1.49 | 7.79 |
| **Inorganic Carbon (%)** | 0.28 | 1.66 |
| **SAR (meq/L)** | 0.23 | 0.29 |
| **pH** | 7.4 | 6.4 |

**Table S3**. Metal concentrations (mg/kg) in the <120 mg/kg TPH and 25,700 mg/kg TPH site soils. The contaminated soil was analyzed in duplicates. The mean concentration and standard deviation are provided in brackets.

| Metal | Background (<120 mg/kg TPH) Soil | Contaminated (25,700 mg/kg TPH) Soil |
| --- | --- | --- |
| **Ag** | <0.25 | <0.25 (0) |
| **Al** | 13,000 | 11,000 (0) |
| **As** | 6.1 | 10.5 (0.50) |
| **B** | <5.0 | <5.0 (0) |
| **Ba** | 66 | 110.0 (0) |
| **Be** | 0.48 | 0.70 (0) |
| **Ca** | 8,300 | 4,550 (150) |
| **Cd** | 0.62 | 0.90 (0) |
| **Co** | 14 | 14.5 (0.50) |
| **Cr** | 37 | 38.0 (4.0) |
| **Cu** | 41 | 74.5 (1.5) |
| **Fe** | 36,000 | 46,000 (1,000) |
| **K** | 1,400 | 1,300 (100) |
| **Mg** | 7,100 | 6,100 (500) |
| **Mn** | 510 | 570 (10.0) |
| **Mo** | 0.66 | 1.10 (0.20) |
| **Na** | 1,100 | 530 (30.0) |
| **Ni** | 28 | 30.0 (1.0) |
| **P** | 560 | 675 (45.0) |
| **Pb** | 13 | 170 (0.0) |
| **S** | 180 | 1015 (185.0) |
| **Sb** | <5.0 | 6.8 (0.1) |
| **Se** | <1.0 | <1.0 |
| **Sn** | <2.0 | 18.5 (0.5) |
| **Sr** | 20 | 22.0 (2.0) |
| **Ti** | 1400 | 1150 (50.0) |
| **Tl** | <1.0 | <1.0 |
| **U** | <5.0 | <5.0 |
| **V** | 99 | 90.5 (1.5) |
| **Zn** | 75 | 205 (5.0) |

At 23°C, the initial surface tension of the Oxoid CM0001 Nutrient Broth culture was 65 mN/m and the final surface tension of the culture was 31 mN/m. A final OD_600_ of 2.81 and cell dry weight of 4.65 g/mL was obtained over the 30-hour period. The initial pH of the medium was 7 and the final culture pH achieved was 6.5. The initial surface tension of the medium was 74.5 mN/m and the final surface tension of the culture was 26 mN/m, indicating successful biosurfactant production. The pH of the medium increased from 6 to 9 following culturing.

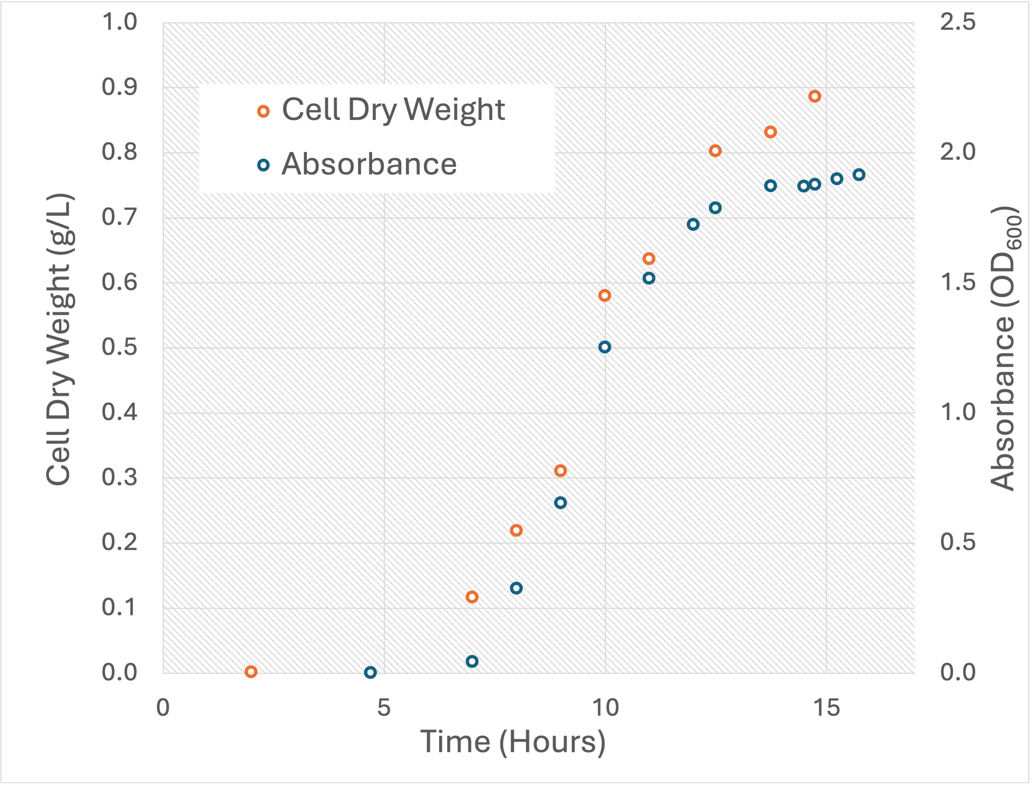

**A**

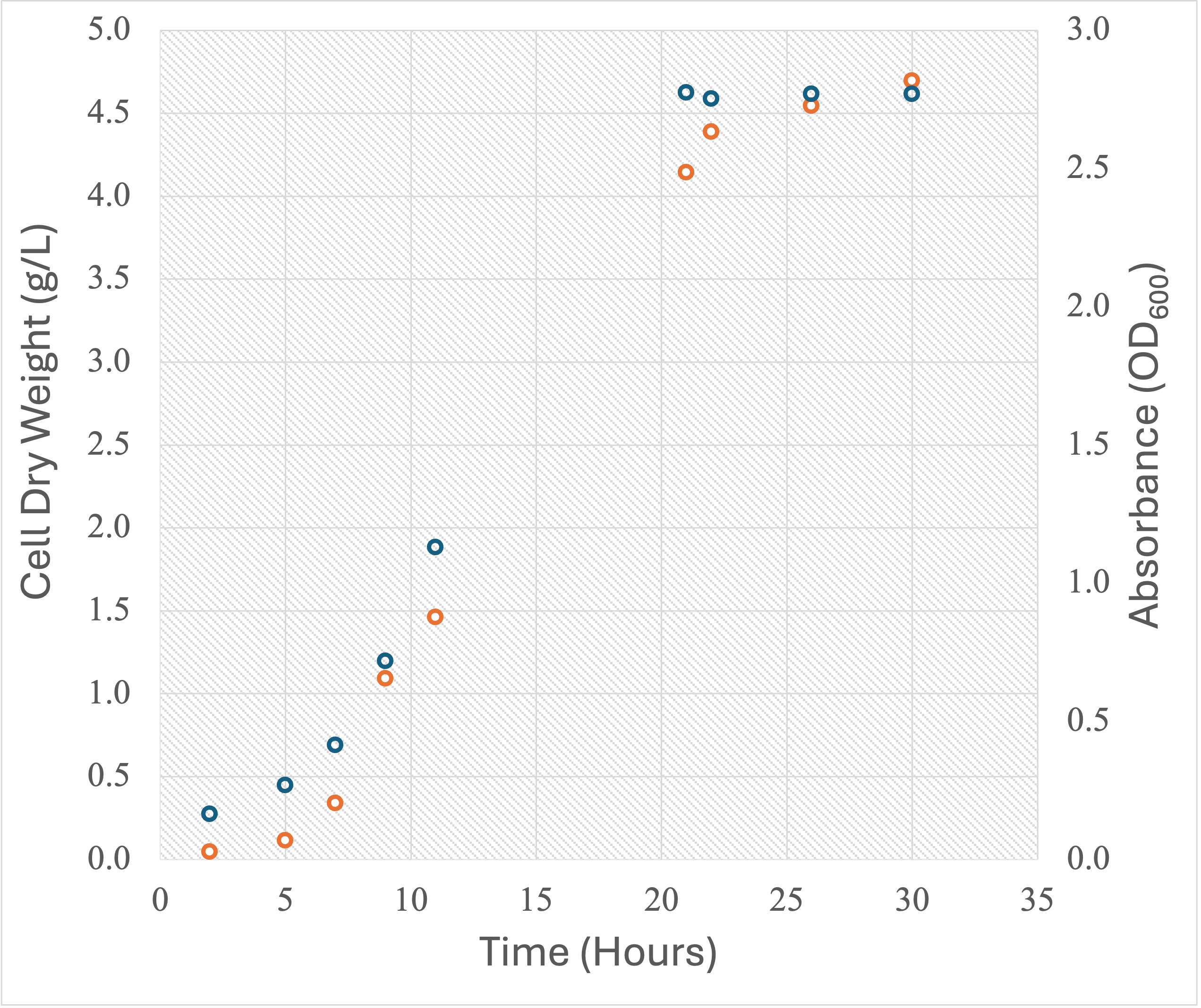

**B**

**Figure S1.** Growth curves of *Bacillus subtilis* ATCC 21332. Cell dry weights (g/L) were measured using a sterile cell filtration unit with Millipore 0.22-micron filter paper. Absorbance readings (OD_600_) were obtained using an Agilent Cary 60 UV-Vis spectrophotometer. **A**. Oxoid CM0001 Nutrient Broth seed culture. **B**. Minerals Salt Medium culture supplemented with 28.3 g/L sucrose. No sucrose remained in the medium following culturing.

Absorbance (OD540)

| Sucrose concentration (g/L) | Standards |
| --- | --- |
| 10 | 0.8609 |
| 20 | 0.9141 |
| 30 | 1.0163 |
| 40 | 1.035 |

| Sucrose concentration (g/L) | Samples |
| --- | --- |
| -114.694 | 0.0894 |
| -110.694 | 0.1142 |
| -104.871 | 0.1503 |
| -94.5323 | 0.2144 |
| -75.6129 | 0.3317 |

**Figure S2**. Concentrations of sucrose in the mineral salt medium following culturing with *Bacillus subtilis* ATCC 21332. Four sucrose standards were tested (blue): 9.4, 18.9, 28.3, and 37.7 g/L sucrose. Remaining sucrose concentrations (orange) was interpolated from the linear curve created using the standards. No sucrose remained following culturing using any of the sucrose concentrations tested.

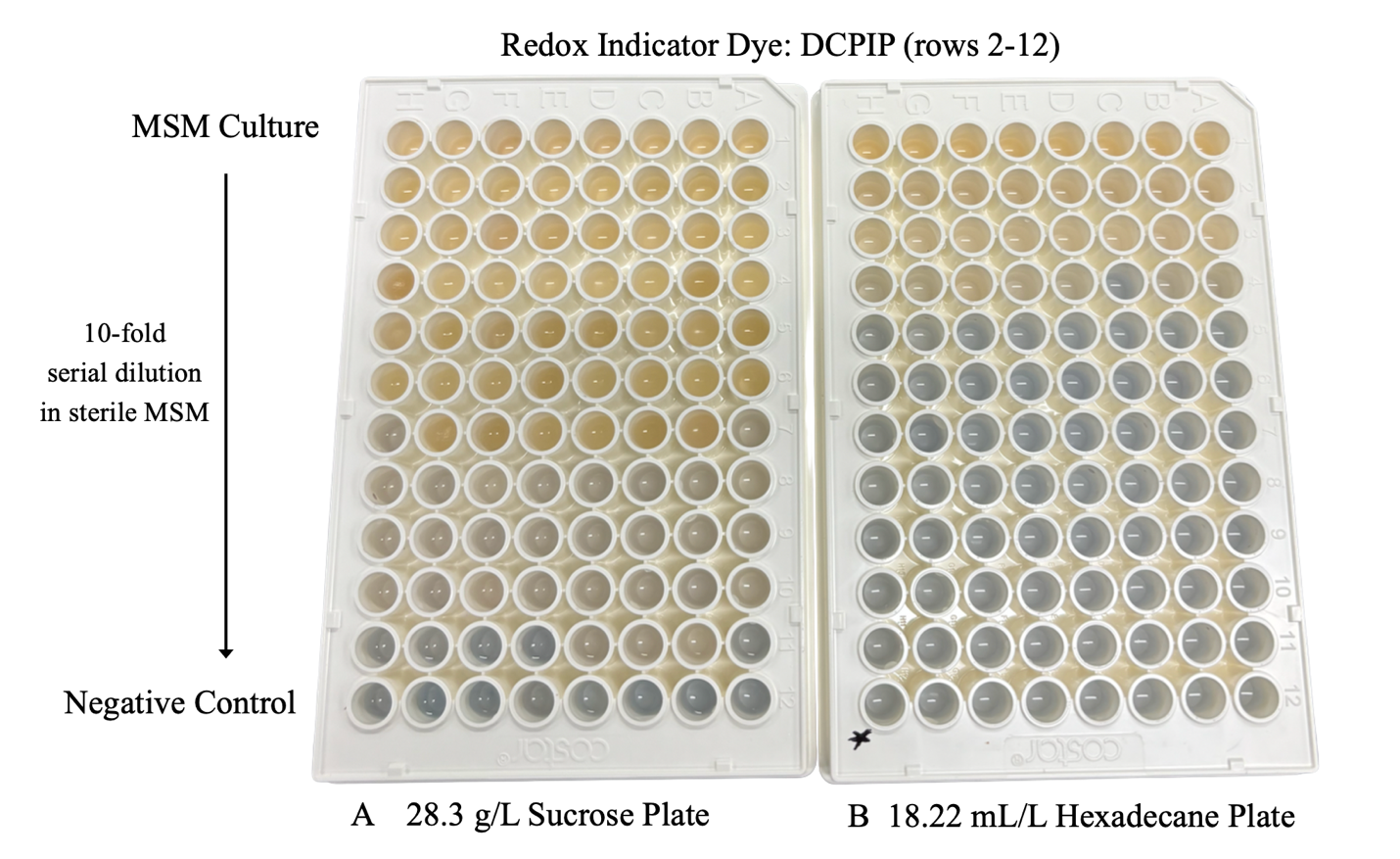

**Figure S3**. Most probable number (MPN) assay of *Bacillus subtilis* ATCC 21332 grown in mineral salt medium supplemented with (**A**) 28.3 g/L sucrose (**B)** or 18.22 mL/L hexadecane. A 10-fold serial dilution was performed using 5 µL of filter-sterilized 2,6-dichlorophenolindophenol (DCPIP) in rows 2-12. Row 1 contained the pure culture whereas row 12 was the negative control. Clear or yellow wells indicate the presence of *B. subtilis*.

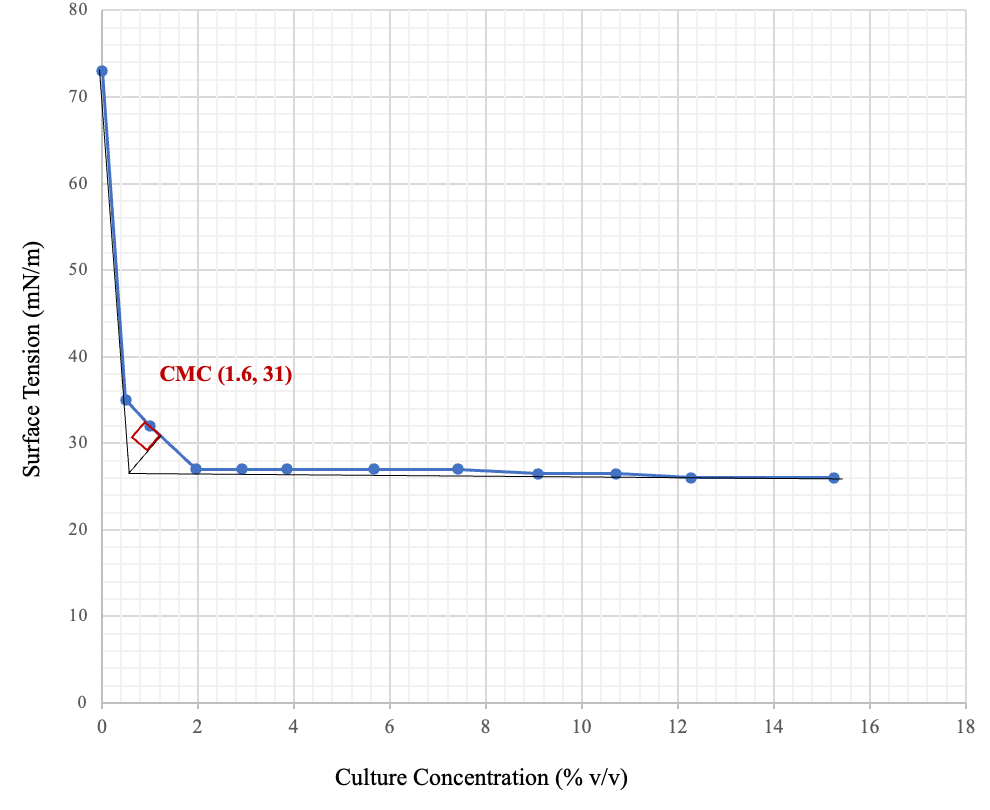

**Figure S4**. Critical micelle concentration of the MSM culture supplemented with 28.3 g/L sucrose. Surface tension (mN/m) was measured using a CSC Precision Dunouy Tensiometer at room temperature (23 °C). Three surface tension readings (*n*=3) were taken at each culture concentration and the mean reading was used.

Critical Threshold

| Log of Quantity (ng/uL) | Critical Threshold (CT) |
| --- | --- |
| 0 | 35.5468 |
| 1 | 29.5609 |
| 1.39794 | 24.11 |
| 1.69897 | 21.85 |
| 1.87506 | 18.93 |

**Figure S5**. Standard curve used to enumerate DNA (ng/µL) during quantitative polymerase chain reactions. The log_10_ of five concentration standards (1, 10, 25, 50, 75 ng/µL) were created from ZymoBIOMICs Microbial Community DNA Standard (2000 ng).

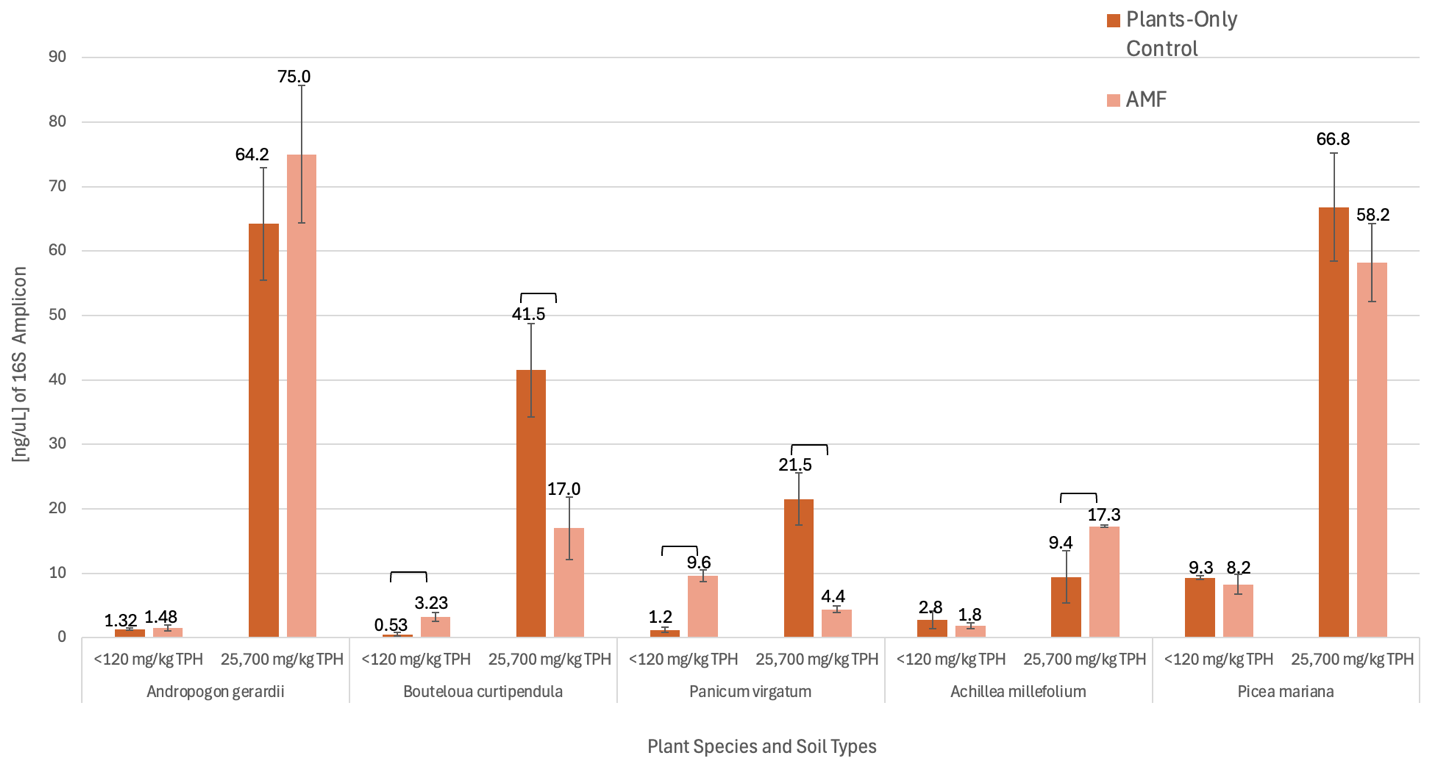

**Figure S6.** Quantification of bacterial DNA in the rhizosphere of the five plant species growing in the <120 mg/kg TPH (background) and 25,700 mg/kg TPH (contaminated) site soils. Bars represent mean 16S amplicon concentration (ng/μL) as determined by qPCR, with error bars indicating standard deviation of the mean. Results are shown for two treatments: Plants-Only Control (dark orange) and AMF (light orange) soil inoculation. Brackets above bars show significant Mann-Whitney U test (*p* < 0.05) pair-wise comparisons between treatments.

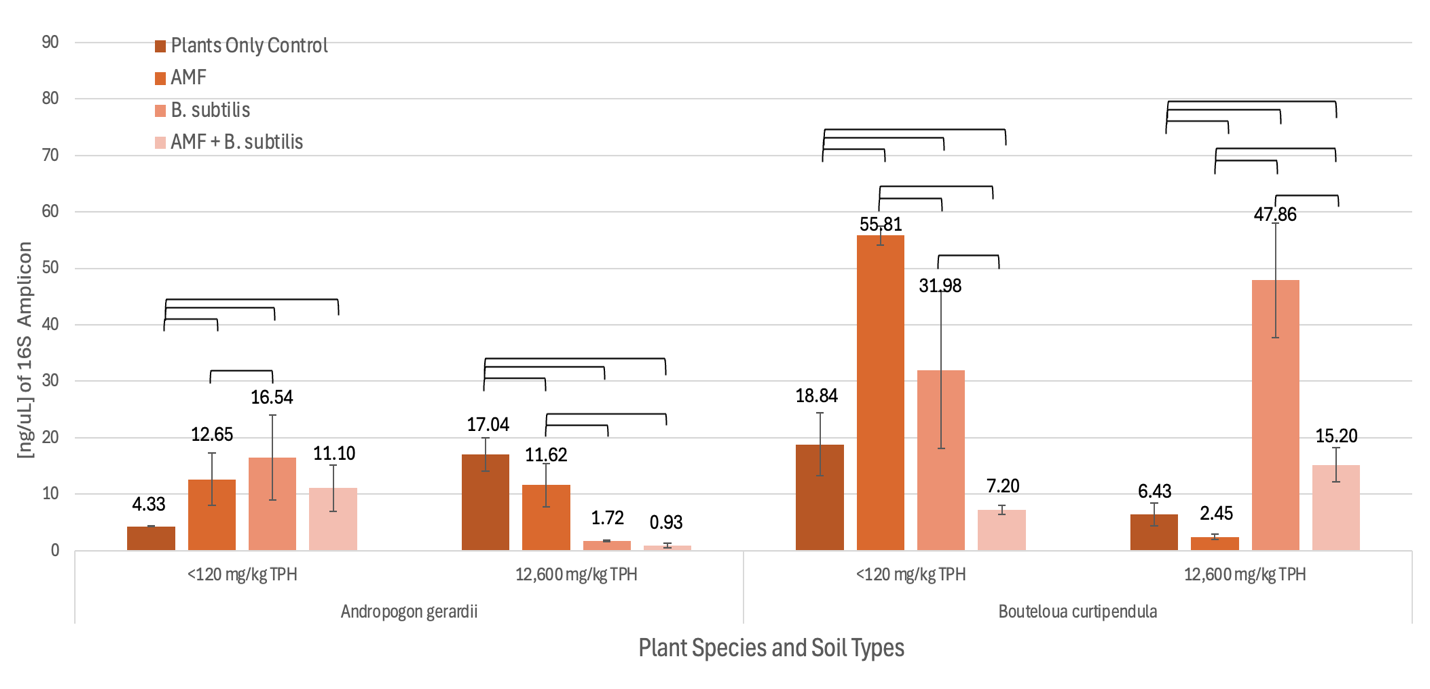

**Figure S7**. Quantification of bacterial DNA in the rhizosphere of *Andropogon gerardii* and *Bouteloua curtipendula* growing in the <120 mg/kg TPH (background) and 12,600 mg/kg TPH (diluted) soils. Bars represent mean 16S amplicon concentration (ng/μL) as determined by qPCR, with error bars indicating standard deviation of the mean. Results are shown for four treatments: Plants-Only Control, AMF, *B. subtilis*, and AMF + *B. subtilis* soil inoculants. Brackets above bars show significant Mann-Whitney U test (*p* < 0.05) pair-wise comparisons between treatments.

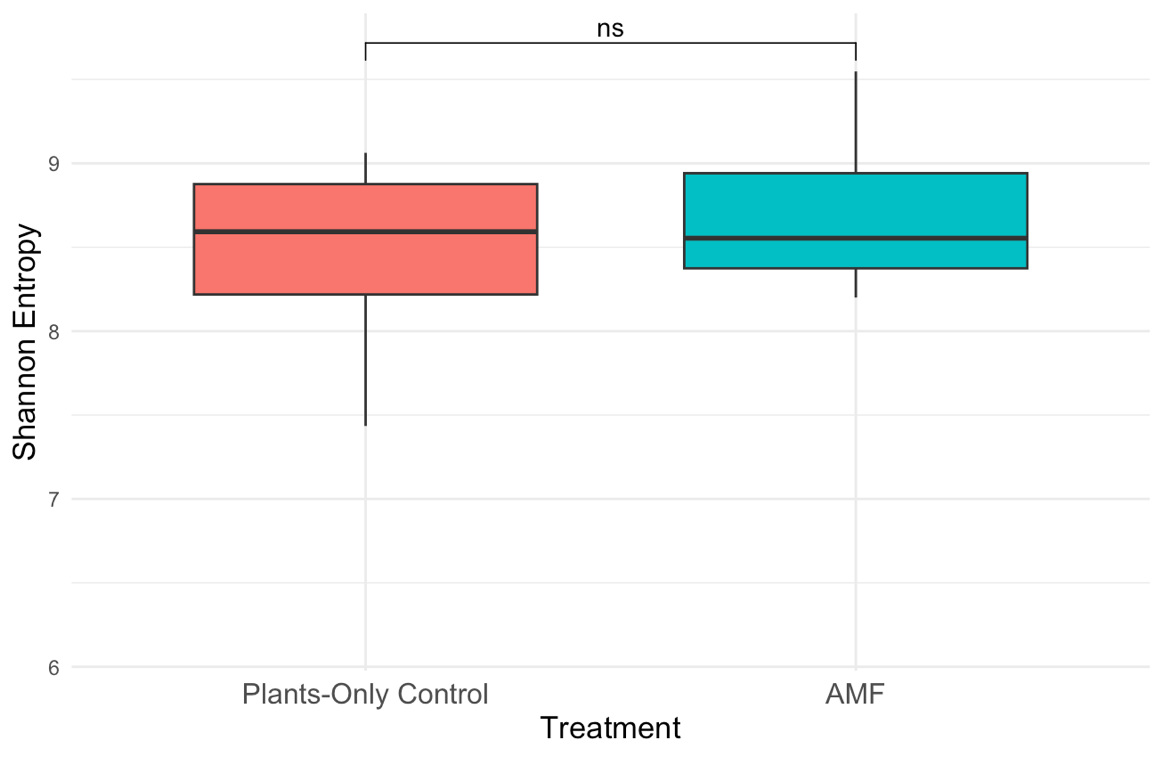

**A**

**B**
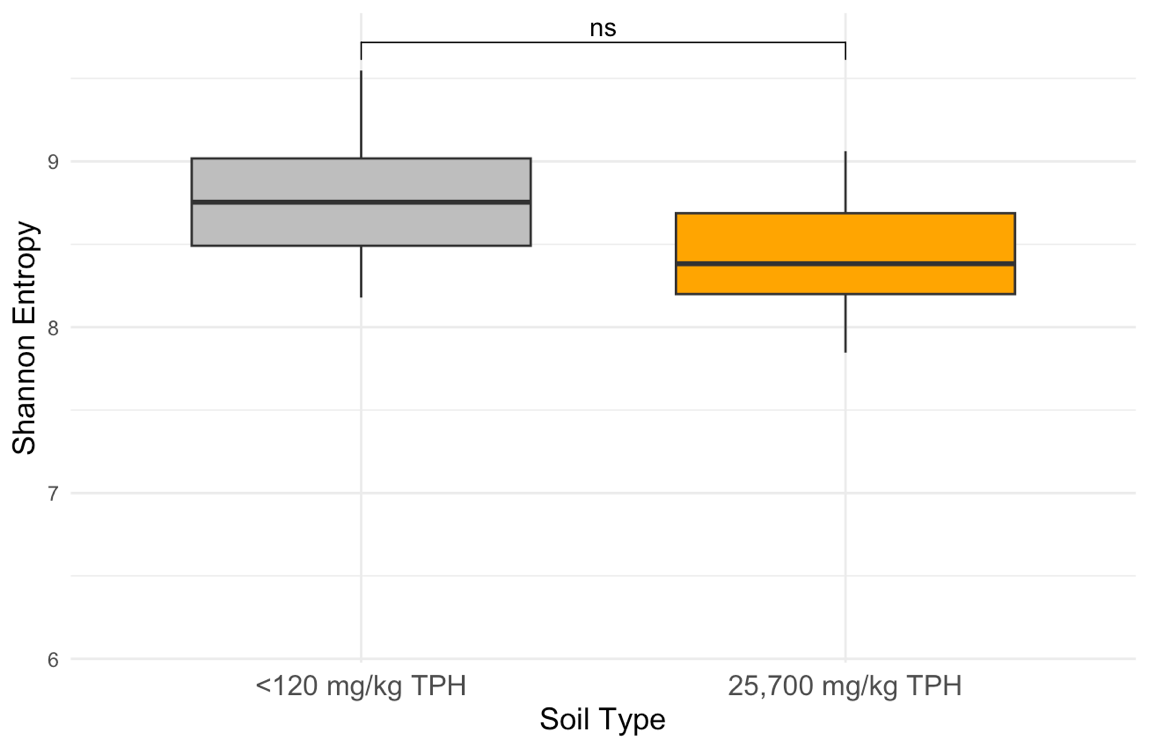

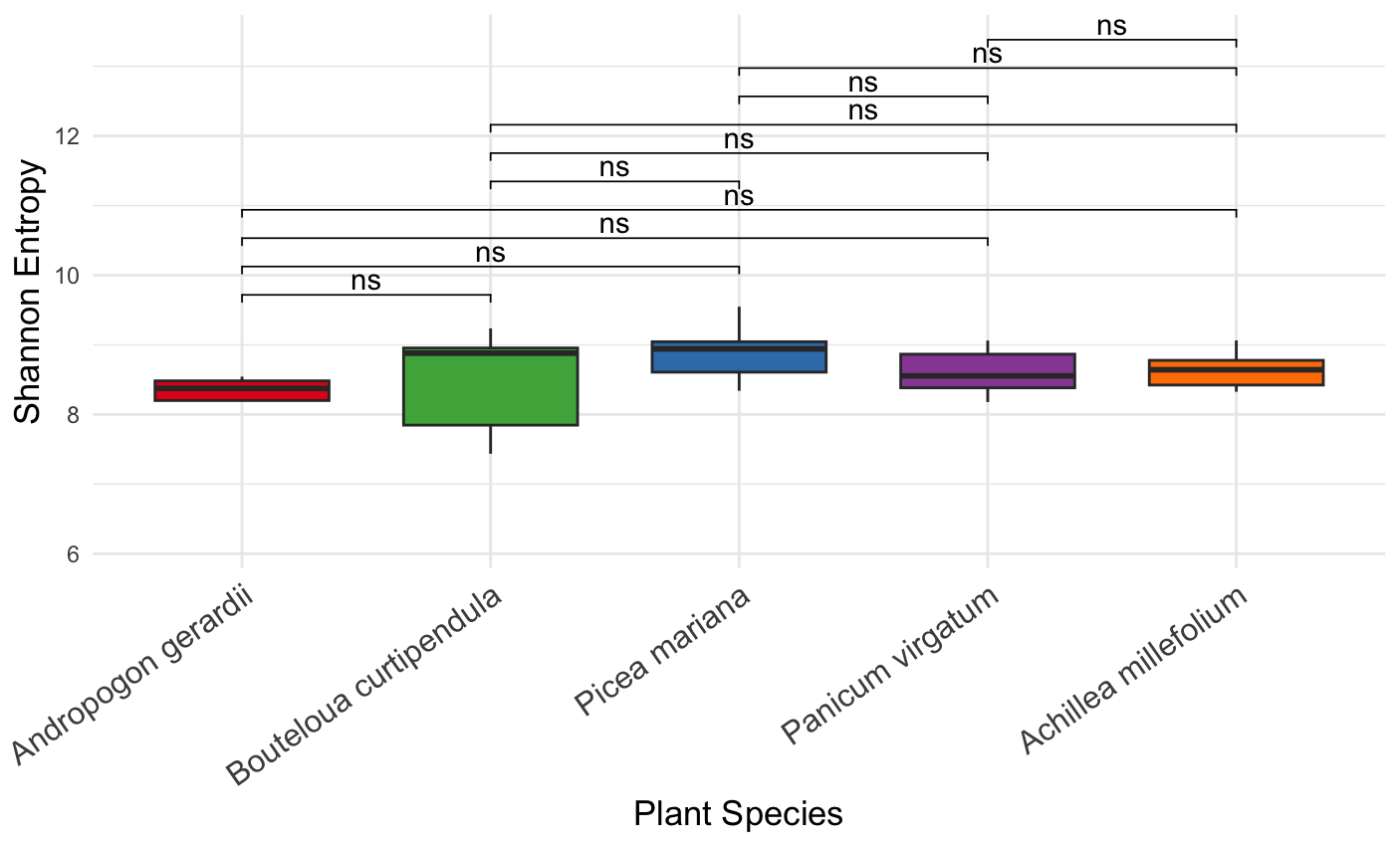

**C**

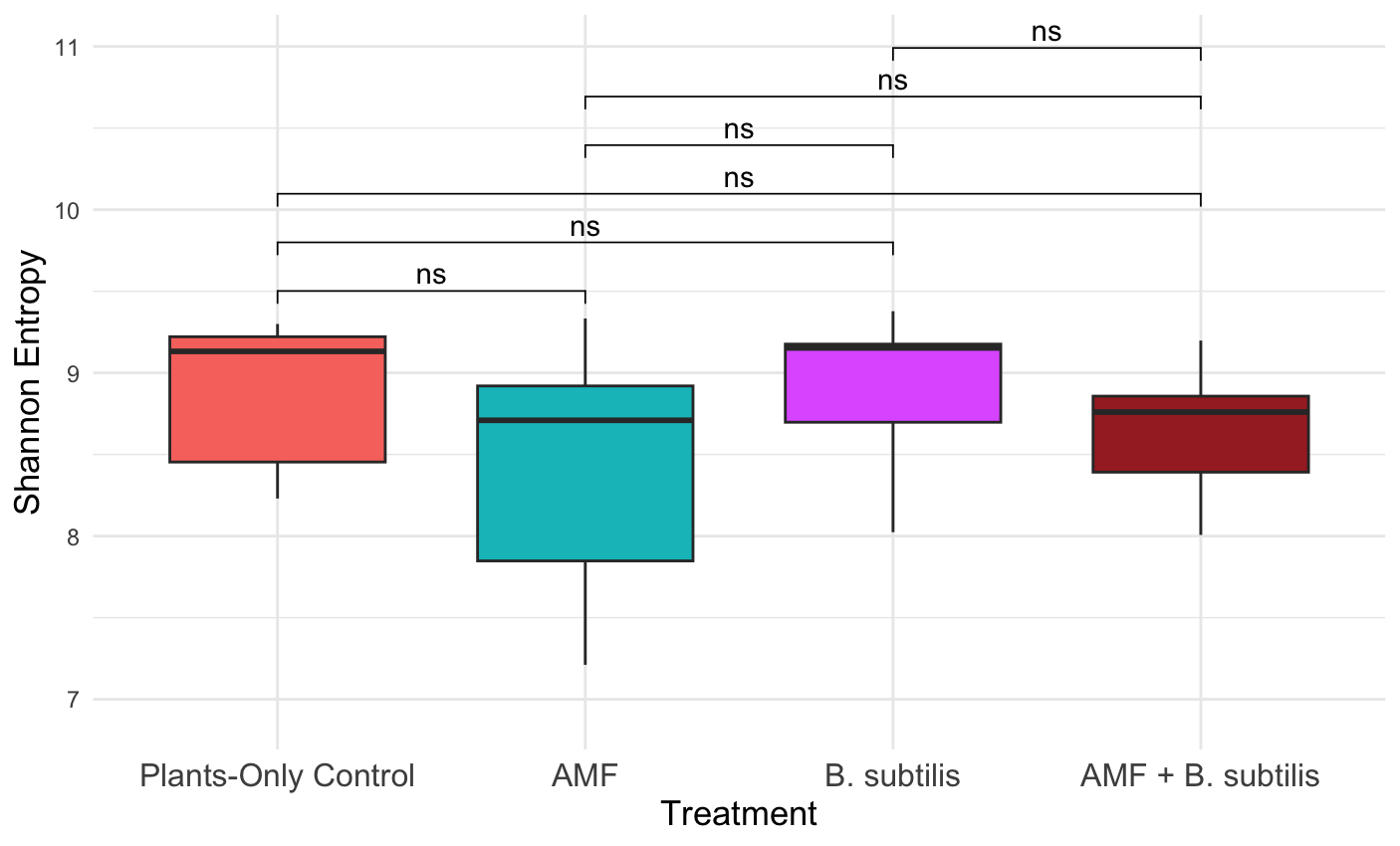

**D**

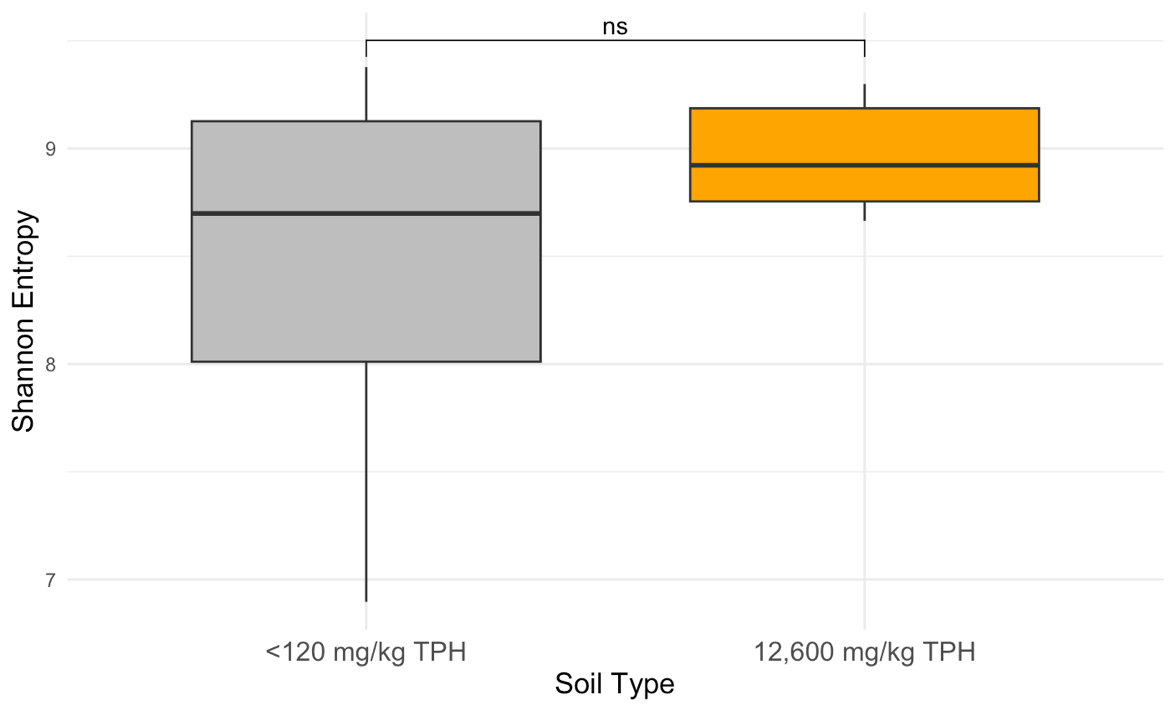

**E**

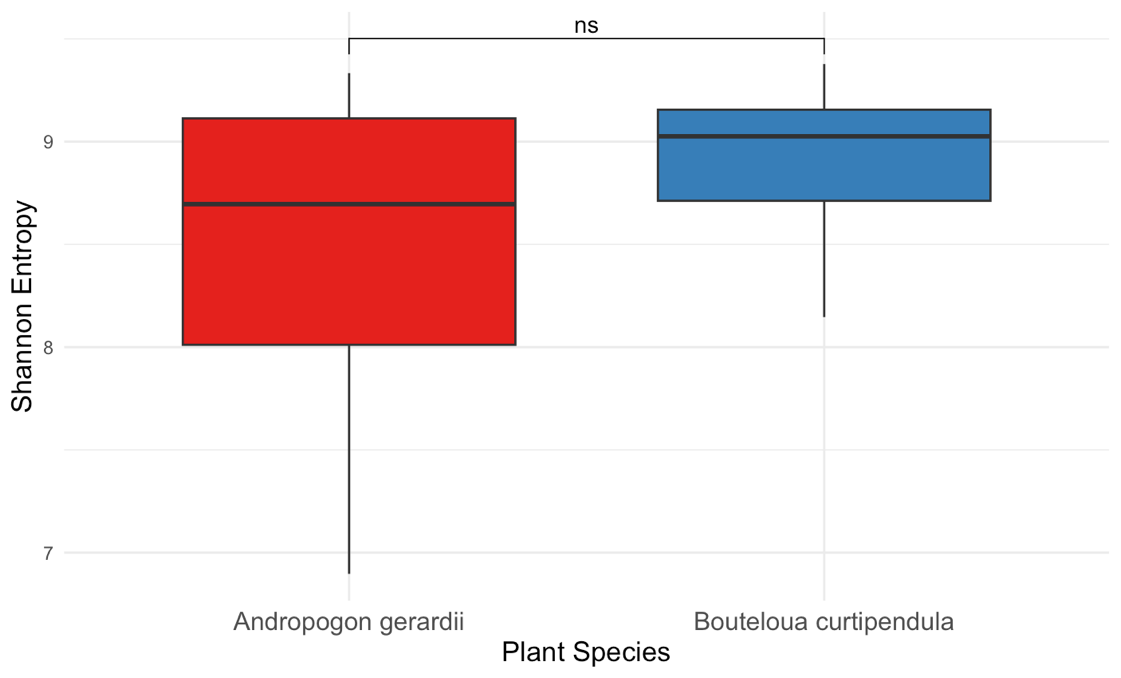

**F**

**Figure S8**. Alpha diversity illustrated through Shannon entropy of the rhizobacteria communities in the northern Ontario soils. Brackets above boxplots show results from Mann-Whitney U test (*p* < 0.05) pair-wise comparisons between treatments, soil types, and plant species. **A.** Treatment (January 2023-2025). **B.** Soil Type (January 2023-2025). **C.** Plant Species (January 2023-2025). **D.** Treatment (June 2023-2025). **E.** Soil Type (June 2023-2025). **F.** Plant Species (June 2023-2025).

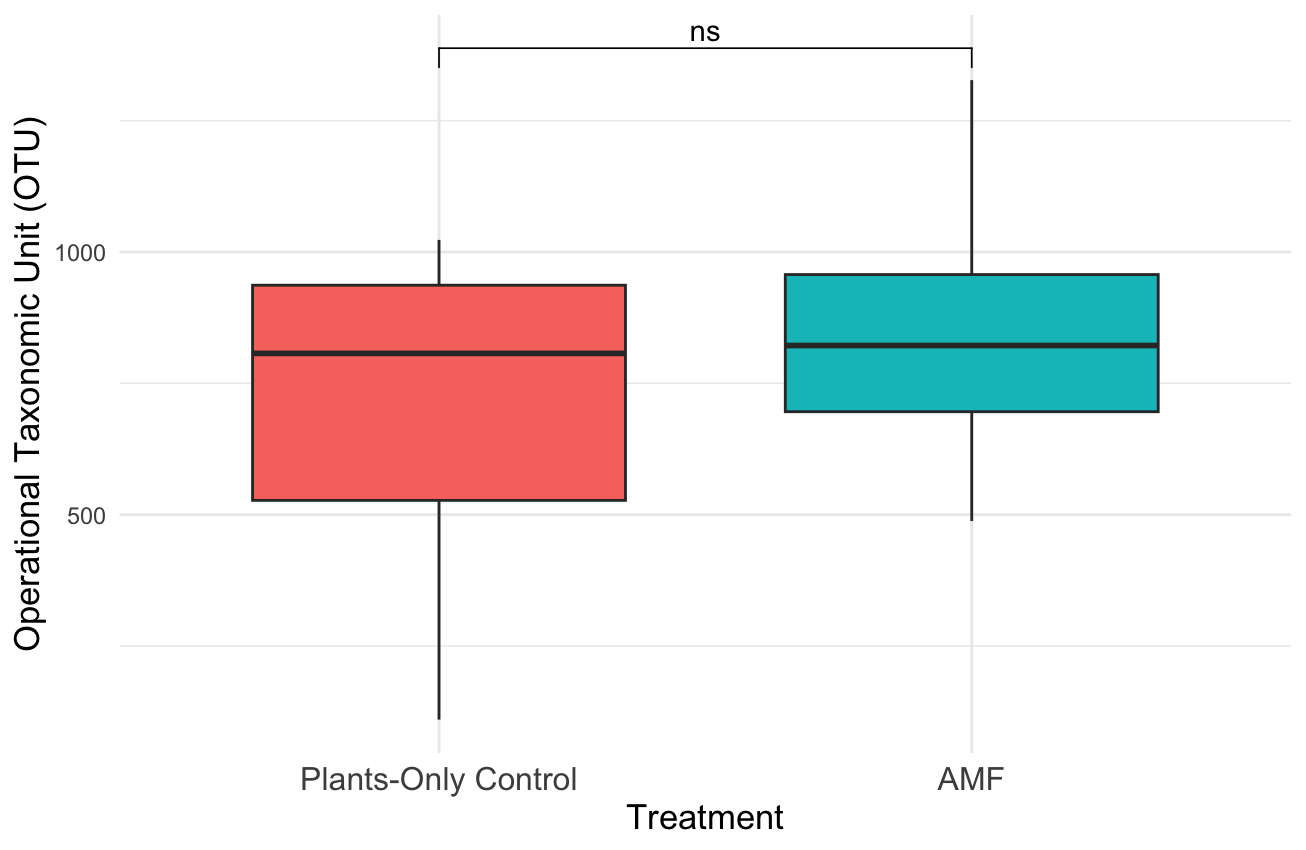

**A**

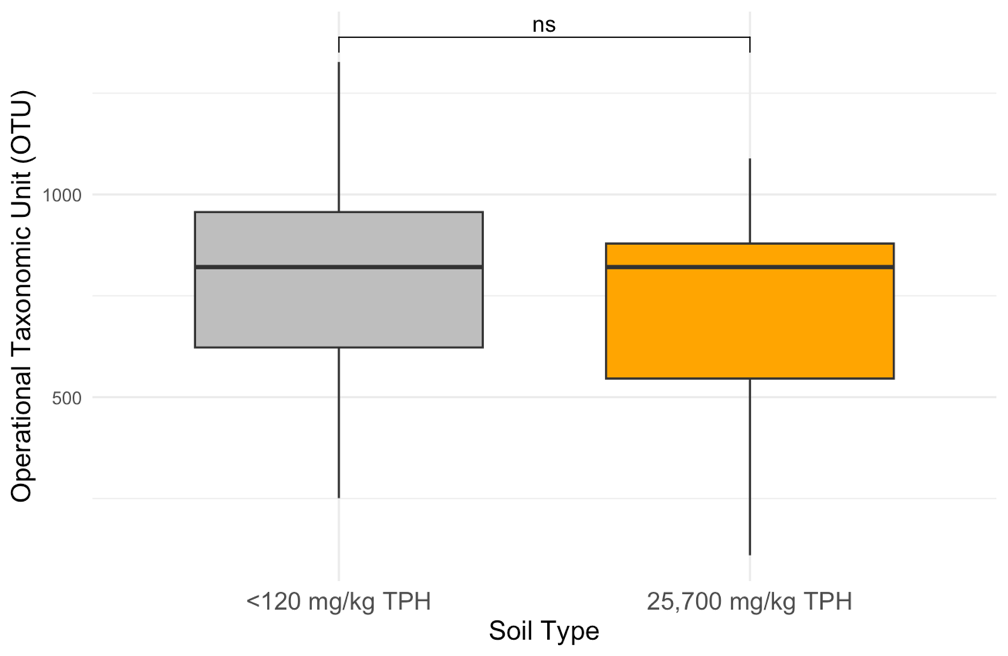

**B**

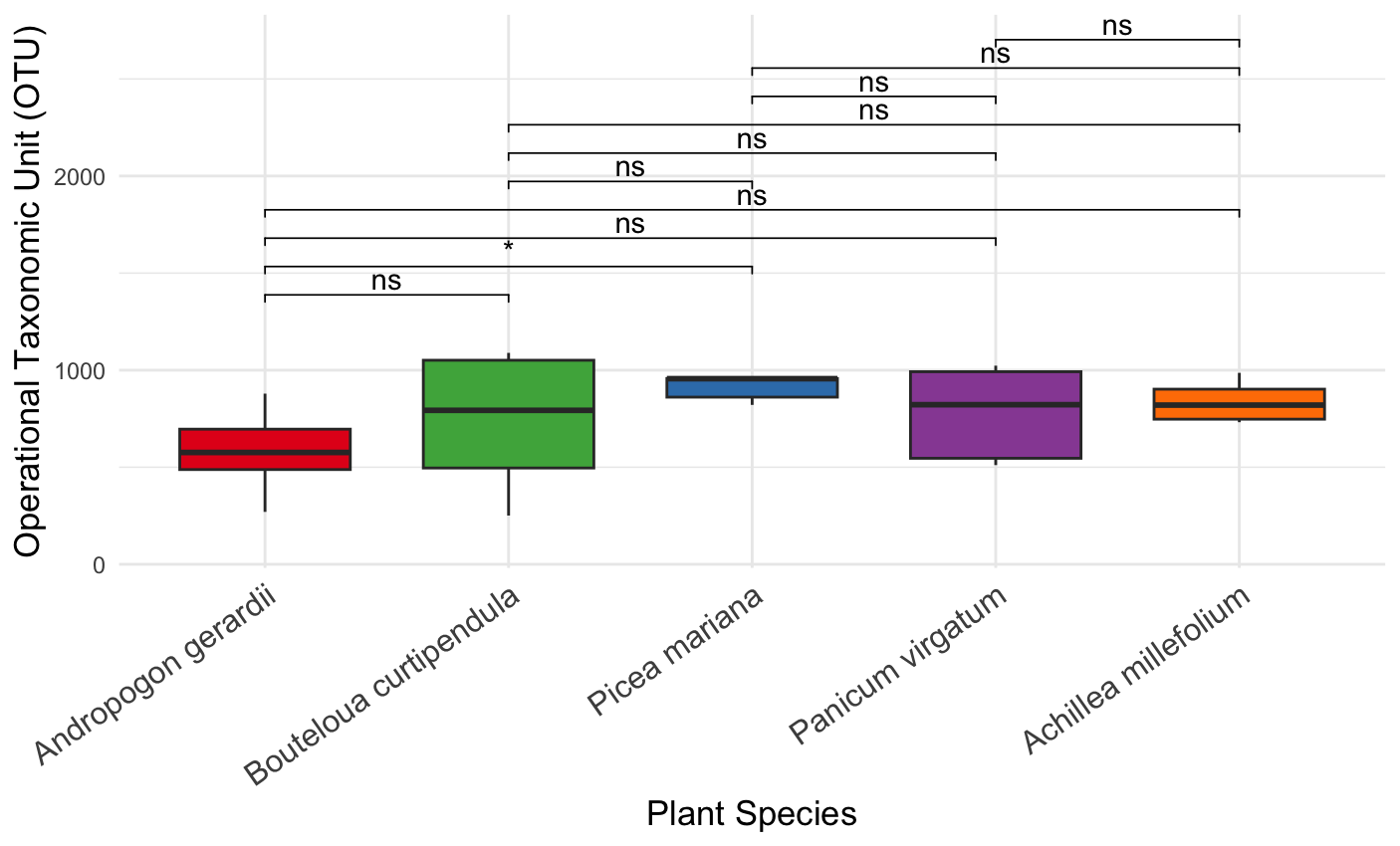

**C**

**D**
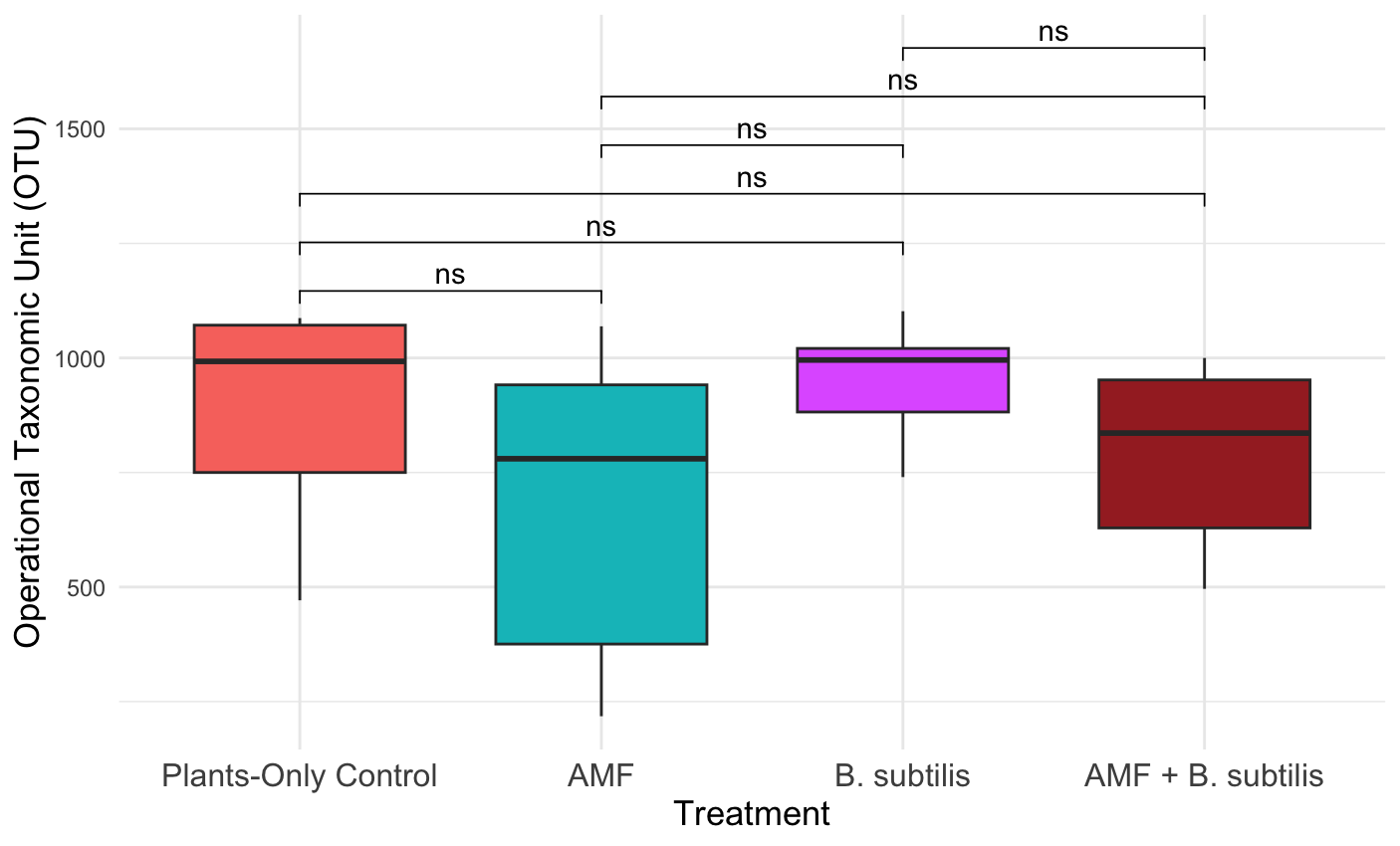

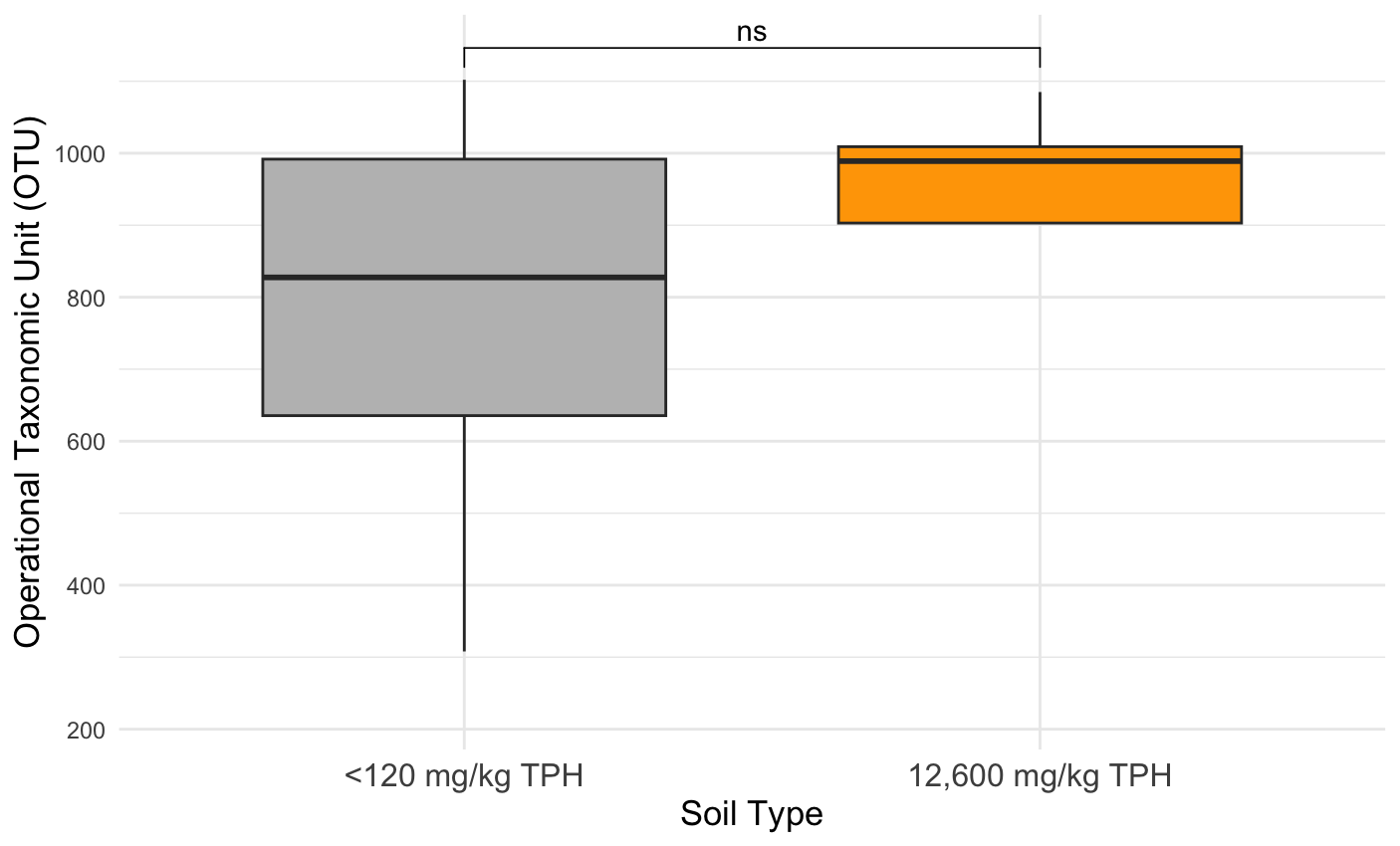

**E**

**F**
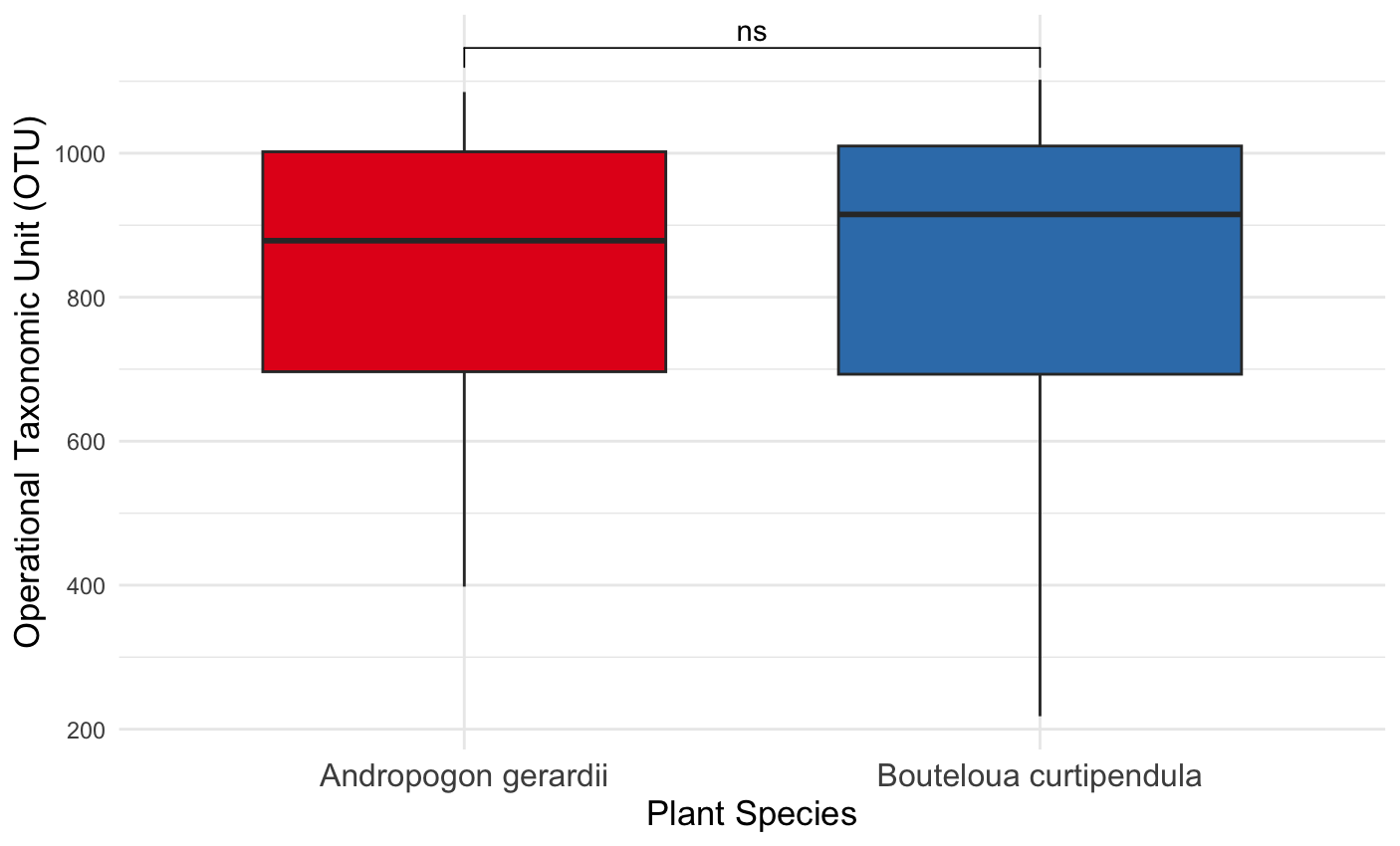

**Figure S9**. Alpha diversity illustrated through Operational Taxonomic Unit (OTU) richness of the rhizobacteria communities in the northern Ontario soils. Brackets above boxplots show results from Mann-Whitney U test (*p* < 0.05) pair-wise comparisons between treatments, soil types, and plant species. **A.** Treatment (January 2023-2025). **B.** Soil Type (January 2023-2025). **C.** Plant Species (January 2023-2025). **D.** Treatment (June 2023-2025). **E.** Soil Type (June 2023-2025). **F.** Plant Species (June 2023-2025).

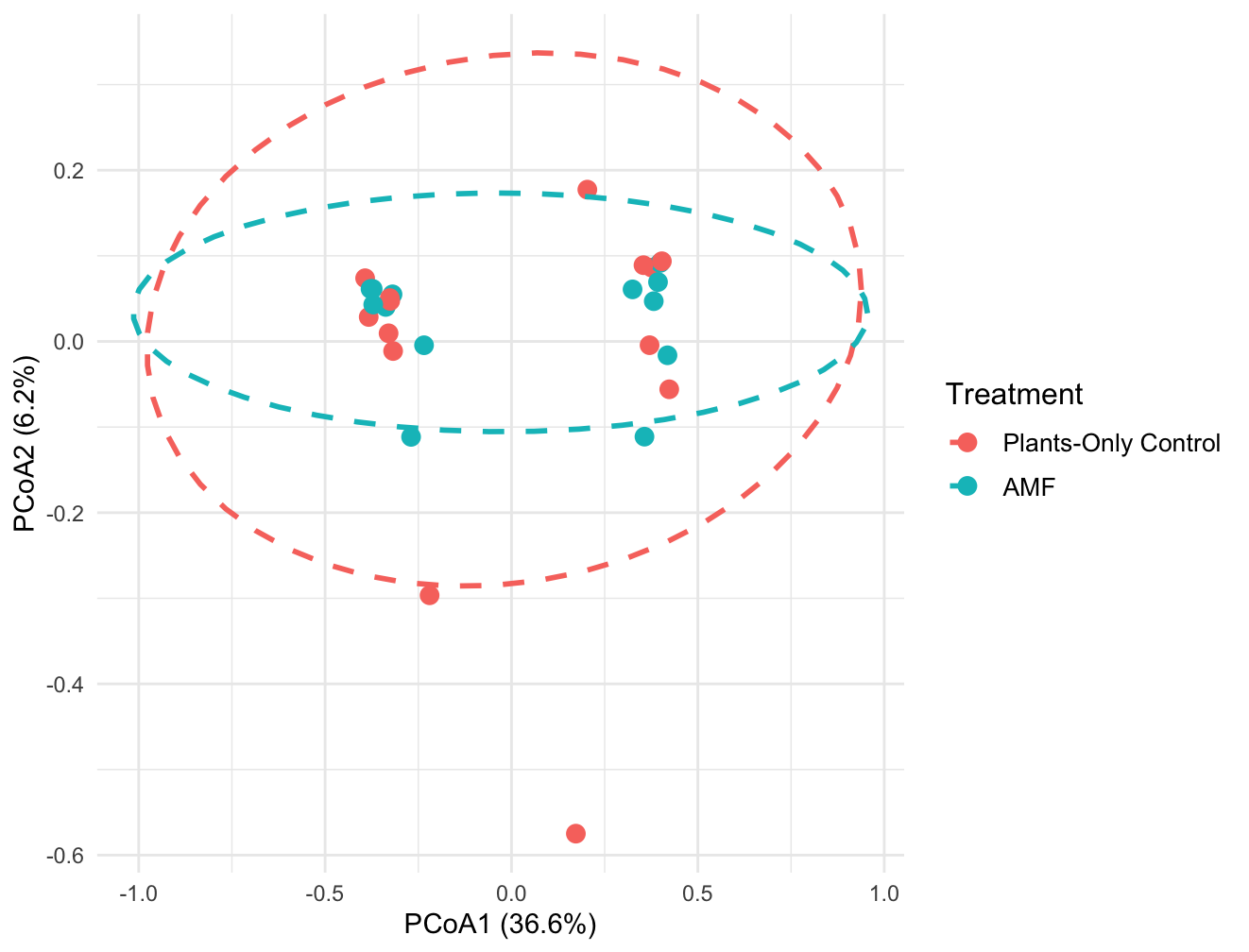

**Figure S10**. Principal coordinates of analysis based on treatment (Plants-Only Control and AMF) during the January 2023-2025 experiments.

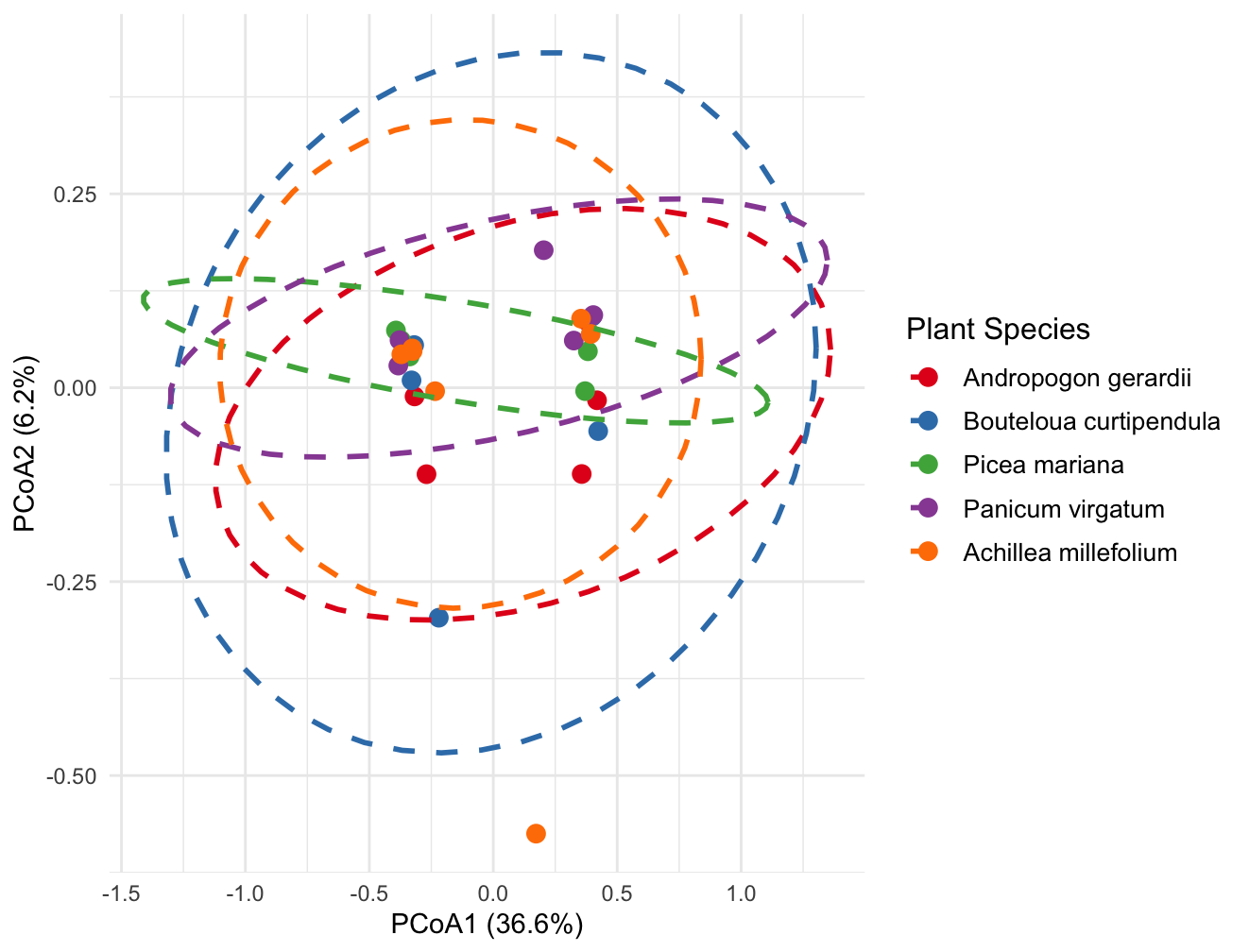

**Figure S11.** Principal coordinates of analysis based on plant species (*Andropogon gerardii*, *Bouteloua curtipendula*, *Picea mariana*, *Panicum virgatum,* and *Achillea millefolium*) during the January 2023-2025 experiments.

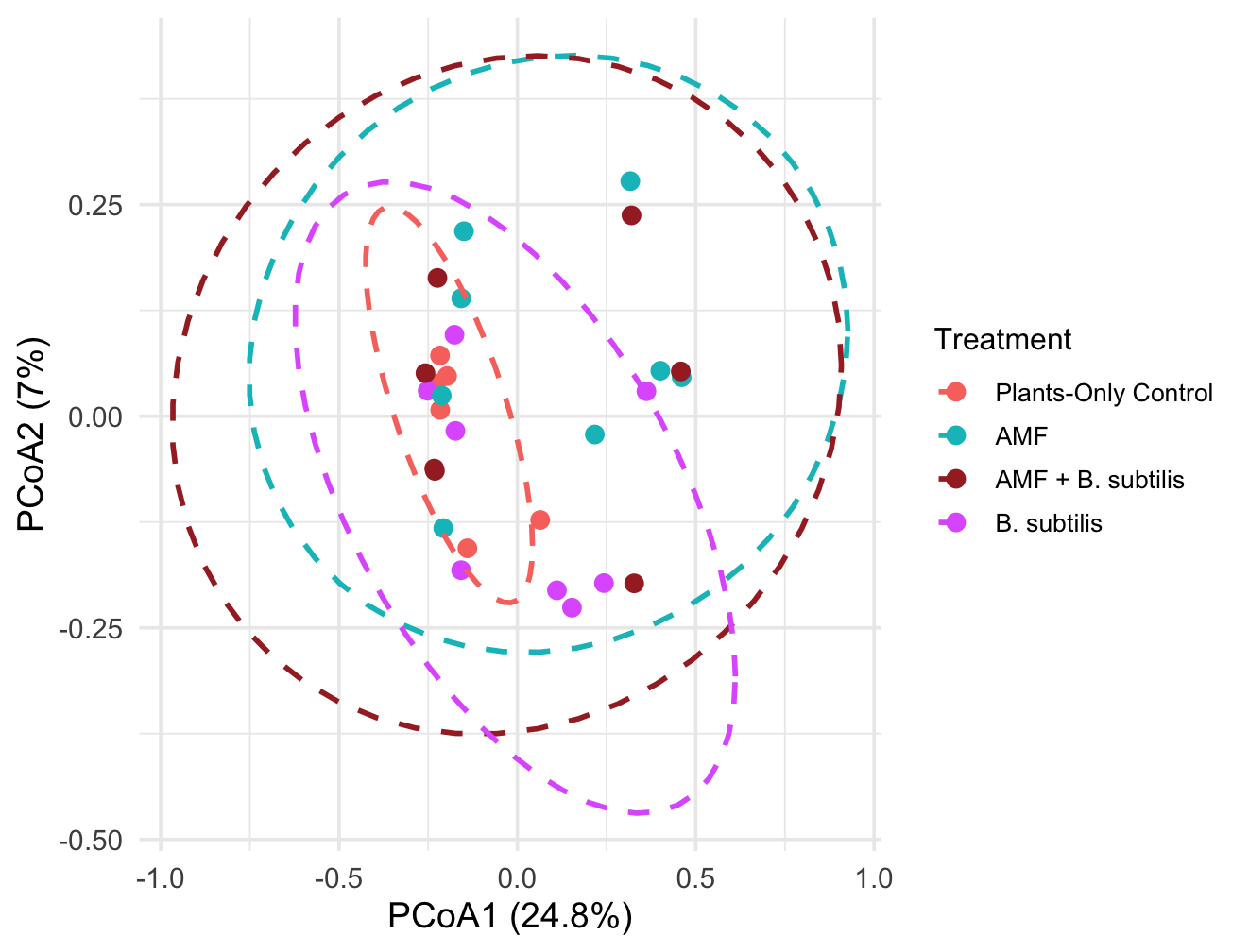

**Figure S12.** Principal coordinates of analysis based on treatment (Plants-Only Control, AMF, AMF + *B. subtilis,* and *B. subtilis*) during the June 2023-2025 experiments.

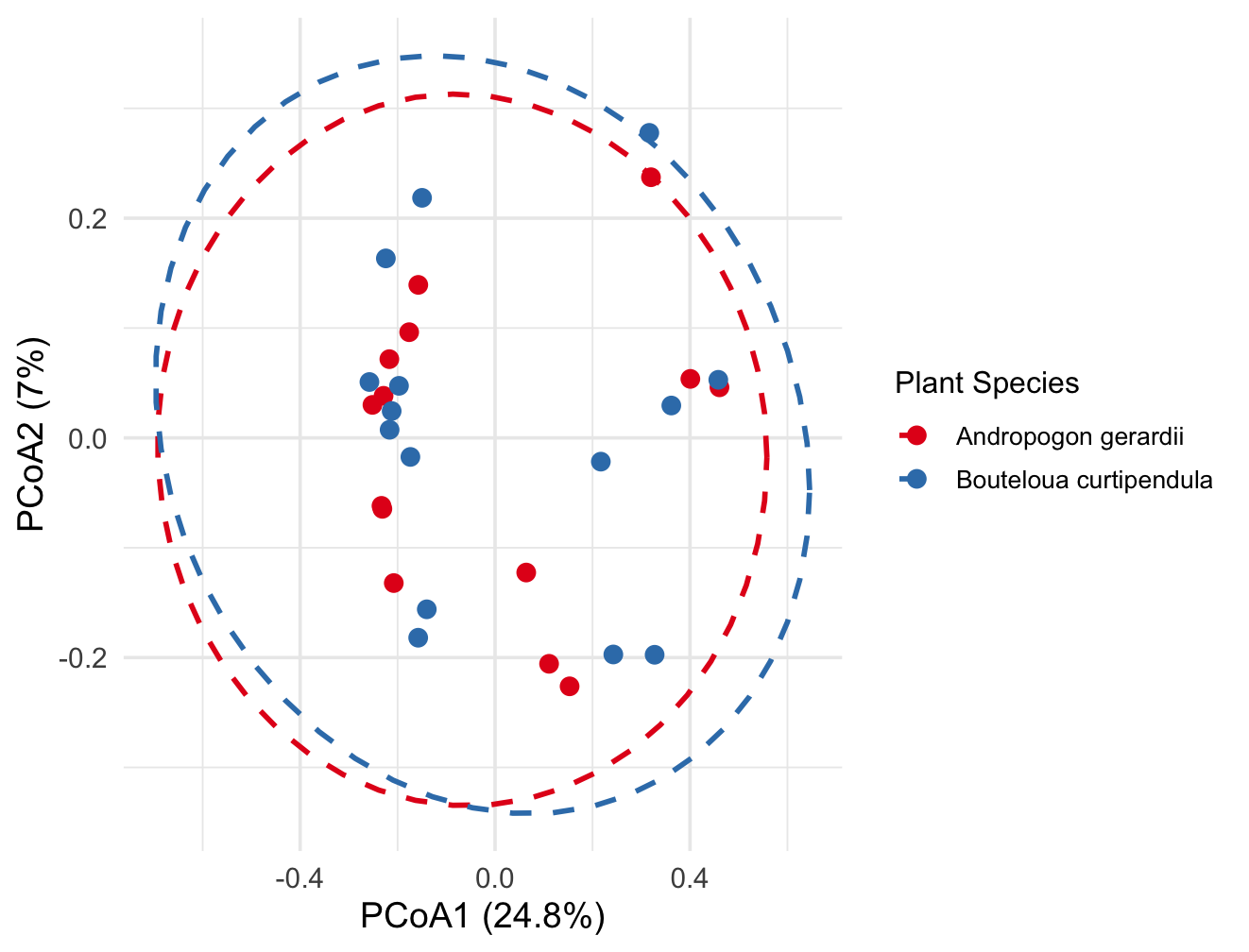

**Figure S13.** Principal coordinates of analysis based on plant species (*Andropogon gerardii* and *Bouteloua curtipendula*) during the June 2023-2025 experiments.

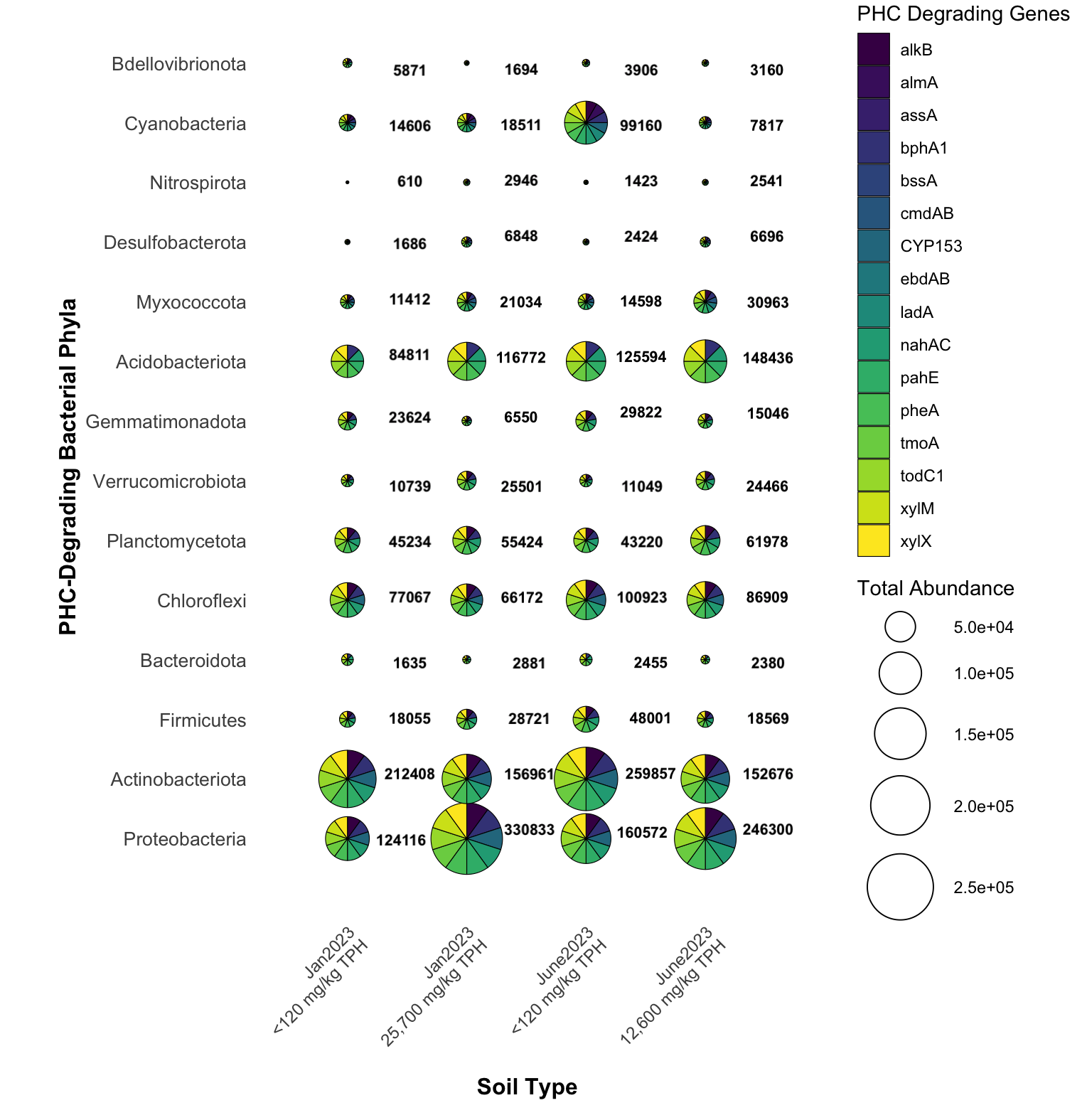

**Figure S14**. Distribution and total abundance counts of PHC-degrading bacterial phyla in the January 2023 - 2025 (“Jan2023”) and June 2023 - 2025 (“June2023”) greenhouse phytoremediation experiments, respectively. Scatterpie bubble charts illustrate phylum-level bacterial communities in the PHC-contaminated (25,700 mg/kg and 12,600 mg/kg TPH) soils and background (<120 mg/kg TPH) soils. Bubble size corresponds to total bacterial abundance within each phylum (numerical values displayed adjacent to bubbles). Pie segments represent the distribution of PHC-degrading genes identified within each bacterial phylum based on functional gene annotation.
